# A PIF- and GUN1-regulated switch in cell axis growth drives cotyledon expansion through tissue-specific cell expansion and division

**DOI:** 10.1101/2025.01.27.635034

**Authors:** Nil Veciana, Guiomar Martín, Elena Monte

## Abstract

Despite its crucial role during seedling deetiolation, cotyledon expansion has been largely overlooked, with hypocotyl elongation favored as the primary phenotypic readout in light signaling research. Here, we investigate how cotyledon expansion is regulated during seedling establishment, and reveal that light-induced cotyledon expansion involves a rapid switch in growth direction - from longitudinal in darkness to transversal upon initial light exposure. Using PIFq- and phyA/phyB-deficient Arabidopsis mutants, we demonstrate that this switch is repressed by PIFs in the dark and promoted by phytochromes under red light. Notably, expansion is antagonistically regulated in the light by GUN1-mediated plastid retrograde signaling. Cotyledon expansion involves rapid epidermis cell expansion, transitioning from rectangular in darkness to characteristic lobed cells in light. Importantly, our findings show that mesophyll extension is driven not only by cell enlargement but also by palisade cell division, consistent with an enrichment of cell cycle-related genes that are antagonistically regulated by the PIF/phy system and retrograde signaling in the cotyledon. Finally, using mutant lines expressing PIF1 and phyB specifically in the epidermis, we establish that epidermal expansion can drive palisade cell growth, while mesophyll cell division is predominantly regulated by light at the tissue-specific level. This study provides a novel framework for investigating cotyledon expansion during seedling deetiolation, incorporating tissue-level regulation. We propose that cotyledons serve as an excellent model for studying morphogenesis and organ geometry, which in plants is governed by directional cell growth.

**SIGNIFICANT STATEMENT:** Despite their critical role in the successful establishment of green photoautotrophic organisms, the mechanisms governing cotyledon expansion during seedling deetiolation remain largely unknown. Our study reveals that the phy/PIF system drives a light-induced shift in the cotyledon growth axis to promote cotyledon growth through tissue-specific cell expansion and division, while GUN1-mediated plastid signals counteract this process when chloroplast biogenesis is impaired.

## INTRODUCTION

Cotyledons are embryonic leaves vital for seedling establishment and the early stages of plant development after germination (Wang *et al*., 2019). In Arabidopsis, they are rich in lipids and serve as nutrient source to fuel seedling development, including the differentiation of the cotyledon embryonic proplastids. If germination occurs in the dark, seedlings undergo skotomorphogenesis or etiolation, exhibiting a fast-growing hypocotyl, a hook to protect the apical meristem, and closed unexpanded cotyledons containing etioplasts. Exposure to light triggers photomorphogenesis or de-etiolation, characterized by an extensive transcriptional reprogramming (Jiao *et al*., 2007) that drives inhibition of hypocotyl growth and chloroplast development in synchrony with the unfolding and expansion of the cotyledons that become the major photosynthetic tissue, a switch to autotrophy critical for seedling survival (Gommers and Monte, 2018).

Photomorphogenesis is initiated by a suite of photoreceptors. Among them, the phytochromes (phy) perceive light of the red and far red wavelengths (600-750nm). The phy family contains five members in Arabidopsis, the photolabile phyA and the photostable phyB-phyE (Ref). Because phyA is the most abundant phytochrome in dark-grown seedlings (Debrieux and Fankhauser, 2010), and phyB is the most abundant and the main contributor in the light (Cantón and Quail, 1999), the concerted action of phyA and phyB regulates the majority of the responses of dark-grown seedlings when first exposed to red light (Tepperman *et al*., 2006). Phys exist in two reversible conformations: the inactive Pr form is synthesized in the cytosol, and absorbs red light to photoconvert to the active Pfr form. Pfr inactivates back to Pr when absorbing far-red light or in prolonged darkness (Ulijasz *et al*., 2010). Upon activation, phy Pfr translocates to the nucleus where it interacts with the basic helix-loop-helix type transcriptional regulators PHYTOCHROME INTERACTING FACTORS (PIFs) (Ni *et al*., 1999). PIFs accumulate in the dark and promote skotomorphogenesis through regulation of about 10% of the genes in the genome (Monte *et al*., 2004; Leivar *et al*., 2008, 2009; Leivar and Monte, 2014). In the light, PIF interaction with active phy Pfr interferes with their DNA binding and induces their rapid phosphorylation and degradation, modifying the gene expression landscape and allowing photomorphogenesis to proceed (Monte *et al*., 2004; Leivar *et al*., 2008, 2009; Park *et al*., 2012). The importance of PIFs as photomorphogenesis repressors during post-germinative development in the dark is manifested by the constitutive photomorphogenic (cop)-like phenotype of the PIF quadruple Arabidopsis mutant *pifq*, which lacks the PIF quartet (PIFq) (PIF1, PIF3, PIF4, and PIF5) and displays short hypocotyl, opened apical hook, and unfolded and expanded cotyledons in the dark (Leivar *et al*., 2009; Shin *et al*., 2009). Light-activated phytochromes also inhibit the COP1-SPA complex, which stabilizes HY5 and other photomorphogenesis-promoting factors (Sheerin *et al*., 2015; Wang *et al*., 2021).

Photomorphogenesis can also be regulated by chloroplast-to-nucleus retrograde signaling (RS), which can converge with phy signaling to antagonistically regulate seedling development (Martin *et al*., 2016). Seedlings treated with lincomycin or norflurazon (inhibitors of chloroplast biogenesis) display closed, unexpanded cotyledons in the light, resembling dark-grown seedlings (Ruckle and Larkin, 2009; Martin *et al*., 2016). RS-induced repression of photomorphogenesis is mediated by GENOMES UNCOUPLED 1 (GUN1) (Ruckle and Larkin, 2009; Martin *et al*., 2016) and is independent of the PIFq (Martin *et al*., 2016) RS and phy signaling converge to regulate *GLK1,* and the GUN1/GLK1 module inhibits the PIF-repressed transcriptional network to suppress cotyledon development when chloroplast integrity is compromised (Martin *et al*., 2016), at least in part through the regulation of BBX16 (Veciana *et al*., 2022).

Cotyledons are structured as stacked combinations of cell layers (Stoynova-Bakalova *et al*., 2004). In the light, the epidermis (upper or adaxial, and lower or abaxial) is characterized by the pavement cells in characteristic jigsaw puzzle arrangement (Lin et al., 2015; Carter et al., 2017) and the embedded stomata (Bean *et al*., 2002; Chen *et al*., 2012; Rovira *et al*., 2024). Between the two epidermises, the mesophyll is composed of two distinct cell layers: the palisade under the adaxial epidermis, consisting of tightly packed column-shaped cylindrical cells (in perpendicular orientation to the cotyledon blade plane) that contain large amounts of chloroplasts and is the main contributor to photosynthesis (Outlaw *et al*., 1976; Gotoh *et al*., 2018); and the spongy between the palisade and the abaxial epidermis, conformed by loosely organized spherical-shaped cells with large air pockets within them and fewer chloroplast, with the primary function of allowing gas flow within the cotyledon/leaf to facilitate photosynthesis (Borsuk *et al*., 2022).

How cotyledon expansion occurs is relatively underexplored. Based on their mutant phenotypes, a role for the phy-PIF module in light-induced cotyledon expansion has been established (Neff and Van Volkenburgh, 1994; Leivar *et al*., 2009; Shin *et al*., 2009; Shi *et al*., 2018), yet how this is achieved is still mostly unknown. Previous work has proposed that is largely a consequence of cell expansion rather than cell division (Neff and Van Volkenburgh, 1994; Stoynova-Bakalova *et al*., 2004), but the relative contribution of each process and the cell layers involved are still open questions in the field. Numerous studies have shown the expansion dynamics of epidermis cells in dark-grown seedlings exposed to light, from the increase in size to the characteristic lobe and jigsaw puzzle formation of the pavement cells (Lin *et al*., 2015; Higaki *et al*., 2016; Carter *et al*., 2017). In contrast, cell division in the epidermis have been mostly conducted around stomatal development, where epidermal cells are produced as division byproducts from stomatal precursors (Houbaert *et al*., 2018; Smit *et al*., 2023). The contribution of this process to the overall cotyledon expansion is unlikely to be significant. In contrast to the epidermis, research on cotyledon mesophyll concerning cotyledon expansion is limited (Stoynova-Bakalova *et al*., 2004).

Here, we examine cotyledon expansion in the dark and address the effect of light during deetiolation, focusing on the contribution of the epidermis and the palisade cell layers. Our results indicate that the phy/PIF system regulates a switch in the axis growth necessary for cotyledon expansion, which is antagonistically regulated by GUN1-mediated RS. Moreover, we show that cotyledon expansion involves not only light-induced cell expansion in the epidermis and the palisade, but also PIF-regulated mesophyll cell division. Using lines expressing epidermis-specific PIF1 and phyB, our data suggest a model whereby the epidermis drives cotyledon expansion supported by tissue-autonomous palisade cell division regulated by the phy/PIF system.

## RESULTS

### Cotyledon expansion during de-etiolation involves a light-induced switch in the growth axis

To assess how cotyledon expansion takes place during deetiolation, we measured cotyledon blade growth in the length and width (at the maximum point) axis, and used cotyledon ratio (length of the cotyledon divided by the width, or L/W) as described (Neff and Van Volkenburgh, 1994) to capture the bidirectional expansion of the cotyledon surface. In 2-day-old dark-grown Arabidopsis seedlings (2dD), cotyledons were longer than wider, displaying an elliptical shape with a ratio value of ∼1.6 (Figure 1a-c). If grown in the dark for longer, the width was maintained unaltered during at least the following 3 days, whereas the length progressively increased, leading to an increase in the ratio in the dark from 1.6 to ∼1.8 (Figure 1b,c). In contrast, if 2dD seedlings were exposed to red light, there was a switch in the preferred growth axis especially apparent after 12h, and this activation of growth in the width axis led to a perfectly round cotyledon (ratio=1 after 48 h of continuous red light) (Figure 1a,d,e) and an increase in the area of ∼7-fold (from 0.13mm^2^ in 2dD to 0.9mm^2^ in 48hR). Together, these observations suggest that light triggers a fast switch in the preferential axis growth of the cotyledon (from length to width) allowing expansion to reach a round cotyledon shape when seedlings are exposed to light.

**Figure 1.**
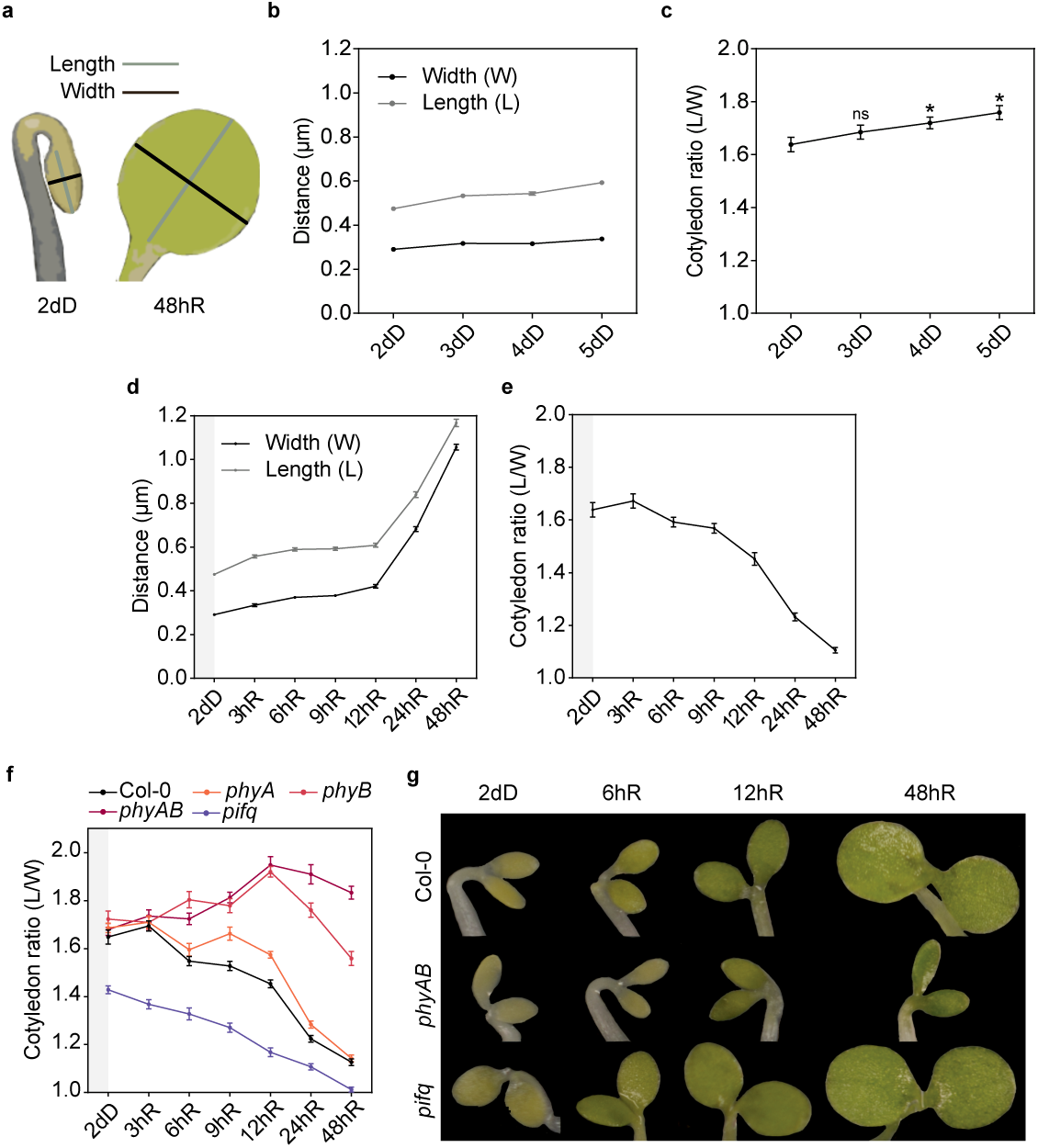
The PHY-PIF module regulates cotyledon expansion and shape during de-etiolation. **(a)** Etiolated (left) and de-etiolated (right) cotyledons with length (gray) and width (black) axes indicated. **(b)** Characterization of cotyledon growth in the two axes in seedlings grown for 2, 3, 4, and 5 days in the dark (dD). **(c)** Quantification of cotyledon ratio (length/width) in WT seedlings grown in (b). Data are means ±SEM of at least 25 seedlings. Statistical differences between mean values at each time point compared to 2dD were analyzed by Student t-test (*P*<0.05) and indicated with an asterisk. ns, non-significant. **(d)** Characterization of cotyledon growth during de-etiolation in the length and width axes. WT seedlings were grown for two days in the dark (2dD) and transferred to continuous red light (R) for 3, 6, 9, 12, 24, and 48 hours. **(e)** Quantification of cotyledon ratio in WT seedlings during de-etiolation. Data are means ±SEM of at least 25 seedlings. **(f)** Quantification of cotyledon ratio in Col-0, *pifq*, *phyA*, *phyB*, and *phyAB* seedlings grown for two days in the dark (2dD) and transferred to continuous red light (R) for 3, 6, 9, 12, 24, and 48 hours. Data are means ±SEM of at least 20 seedlings. **(g)** Visual phenotypes of representative Col-0, *phyAB*, and *pifq* cotyledons during de-etiolation at selected time points.

### The PHY-PIF module promotes the light-induced axis growth switch

To start to understand how expansion in the two axis is regulated, we next performed a de-etiolation experiment using *phyA*, *phyB*, *phyAB*, and *pifq*. In 2dD seedlings, none of the three *phy* mutant lines showed any difference in cotyledon ratio compared to Col-0 (WT) (∼1.6). In contrast, *pifq* displayed a lower ratio of ∼1.4, as expected given their expanded cotyledon phenotype in the dark (Figure 1f) (Leivar *et al*., 2009; Shin *et al*., 2009). Upon red-light light exposure, *phyA* mutants exhibited delayed cotyledon expansion during the first hours (9-12 h) but eventually resumed growth until cotyledon ratio reached WT-like levels after 48 hours of red-light treatment (∼1.1). Remarkably, in *phyB* and *phyAB* mutants, cotyledons not only did not reduce their length/width cotyledon ratio, but the ratio increased in red light (Figure 1f,g) (∼1.9 after 24h of red), revealing a lack of the width-growth priority and a continuous growth in the length axis, mimicking the behavior observed in dark cotyledons (Figure 1c). This was not sustained in *phyB –* eventually expanded after 48 hours – but was maintained in *phyAB* after two days of red light (Figure 1f,g), in accordance with phyA and phyB acting in concert during deetiolation (Tepperman *et al*., 2006). These results support the idea that cotyledon expansion requires a light-induced switch in the preferential axis growth of the cotyledon, and indicate that this switch is repressed by PIFs in the dark and is activated by phy signaling during the first hours of light.

### Convergence of phy/PIF and RS in the regulation of cotyledon expansion

GUN1-mediated inhibition of cotyledon expansion upon chloroplast disruption in continuous light has been described before (Ruckle and Larkin, 2009; Martin *et al*., 2016). However, whether GUN1-mediated RS also affects cotyledon expansion in the dark and/or in dark-grown seedlings when first exposed to light remains to be determined. We first evaluated how lincomycin affected cotyledon ratio in 2dD WT seedlings. Compared to non-treated seedlings, ratio was similar (∼1.6) (Figure 2a,b). However, upon red light exposure for 24h, whereas non-treated cotyledons expanded to a ratio of ∼1.35, lincomycin inhibited WT cotyledon expansion and ratio remained at ∼1.6 (Figure 2a,b). Similarly to WT, *gun1* cotyledon expansion was unaffected by lincomycin in the dark. In contrast, upon transfer to red light, *gun1* cotyledons expanded even in the presence of lincomycin, and exhibited similar reduced ratios compared to dark-grown *gun1* in non-treated (∼1.25) and treated (∼1.4) conditions (Figure 2a,b). These results are in agreement with the previously described GUN1 accumulation in cotyledons of dark- and light-grown as well as in deetiolating seedlings (Wu *et al*., 2018; Hernández-Verdeja *et al*., 2022), and together they show that chloroplast disruption suppress cotyledon expansion in dark-grown seedlings during the first hours of red light exposure, maintaining etiolated-like cotyledons. This suppression of photomorphogenesis by RS is GUN1-mediated, and indicates that the previously described antagonistic convergence of light and RS shown under continuous light conditions (Martin *et al*., 2016) also regulates cotyledon development during dark-to-light deetiolation. Interestingly, cotyledon expansion can proceed with a damaged chloroplast in the *gun1* mutant, suggesting that expansion can be fueled by the remaining seed energy in the cotyledons, and indicating that lack of expansion in lincomycin-treated WT is likely not due to a lack of chloroplast-derived nutrients but rather a consequence of active chloroplast-to-nucleus signaling mediated by GUN1.

**Figure 2.**
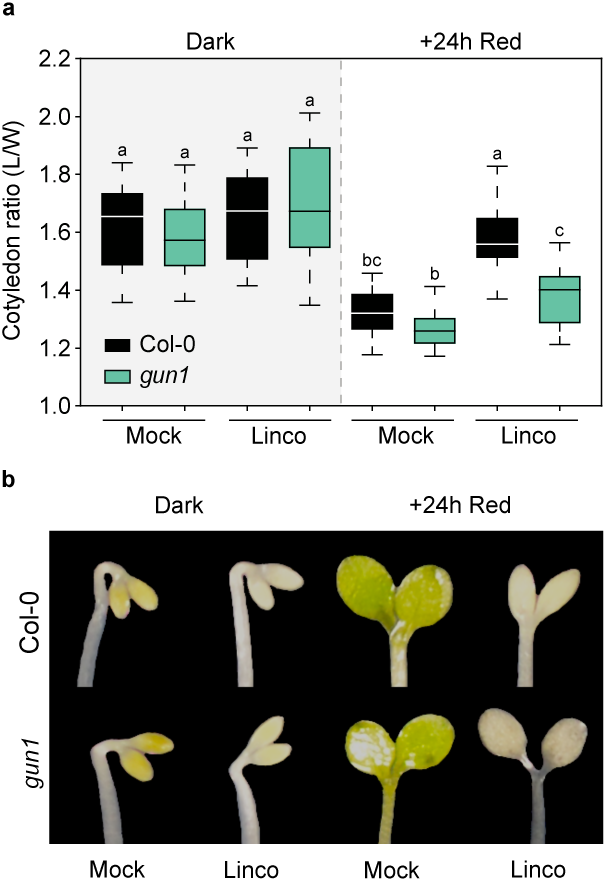
GUN1-mediated retrograde signaling regulates cotyledon expansion and shape during de-etiolation. **(a)** Cotyledon ratio (length/width) quantification of WT Col-0 and *gun1* seedlings grown for two days in the dark and transferred to red light for 24 hours, without (mock) or with (Linco) lincomycin. Data in boxplots indicate the first quartile, median, and third quartile of n≥20 seedlings. Whiskers indicate 5 to 95 percentile. Letters denote the statistically significant differences using two-way ANOVA followed by post hoc Tukey’s test (*P*<0.05). **(b)** Visual phenotypes of representative Col-0 and *gun1* cotyledons of seedling grown as in (a).

### The cotyledon expression network is antagonistically regulated by phytochrome and retrograde signals with an enrichment in cell division-related genes

To start to understand the underlying transcriptional network involved in cotyledon expansion in response to light and retrograde signaling, we first took advantage of a previous report that described light-regulated genes in dark-grown seedlings exposed to light separately in cotyledon and hypocotyl, with a 2-fold change threshold (Sun *et al*., 2016). We reasoned that significant candidate genes would likely belong to categories encompassing light regulated genes in cotyledon that are not affected in hypocotyl, and genes that are expressed in opposite direction in these two organs given the promoting and inhibiting effect of light in cotyledon and hypocotyl cell expansion, respectively (Bou-Torrent *et al*., 2008; Martín and Duque, 2021). Accordingly, we defined two groups (Figure 3a): light up-regulated in cotyledons (Light UP Cot) (2122 genes in total, 1810 cotyledon specific and 312 also down-regulated in hypocotyls), and light down-regulated in cotyledons (Light DOWN Cot) (1781 genes in total, 1640 cotyledon specific and 141 also up-regulated in hypocotyls). In contrast, we considered two additional groups containing light up- or light down-regulated genes in both cotyledons and hypocotyls (Light UP Cot+Hyp, 1178 genes; Light DOWN Cot+Hyp, 679 genes) (Figure 3a) (Supplemental Table 1) as less likely to be involved in the regulation of cotyledon expansion. We next examined the regulation of the “Light UP Cot” and “Light DOWN Cot” gene subsets in *pifq* in the dark, analyzing cotyledon-specific data from the same report (Sun *et al*., 2016). Our results indicate that the light-regulated genes in cotyledon correspond predominantly to PIF-regulated genes and show a similar trend in expression in *pifq* in the dark (Figure 3b), in accordance with their regulation by the phy/PIF system as previously concluded (Sun *et al*., 2016). Finally, to evaluate their regulation by RS, we compared their expression upon treatment with norflurazon (NF), assessing data from Zhao *et al*., (2019). The light-regulated molecular phenotype in both the “Light UP Cot” and “Light DOWN Cot” gene subsets was strongly reversed when RS was activated (Figure 3b). Importantly, this reversion was partially dependent on GUN1 (Figure 3b) (Zhao *et al*., 2019).

**Figure 3.**
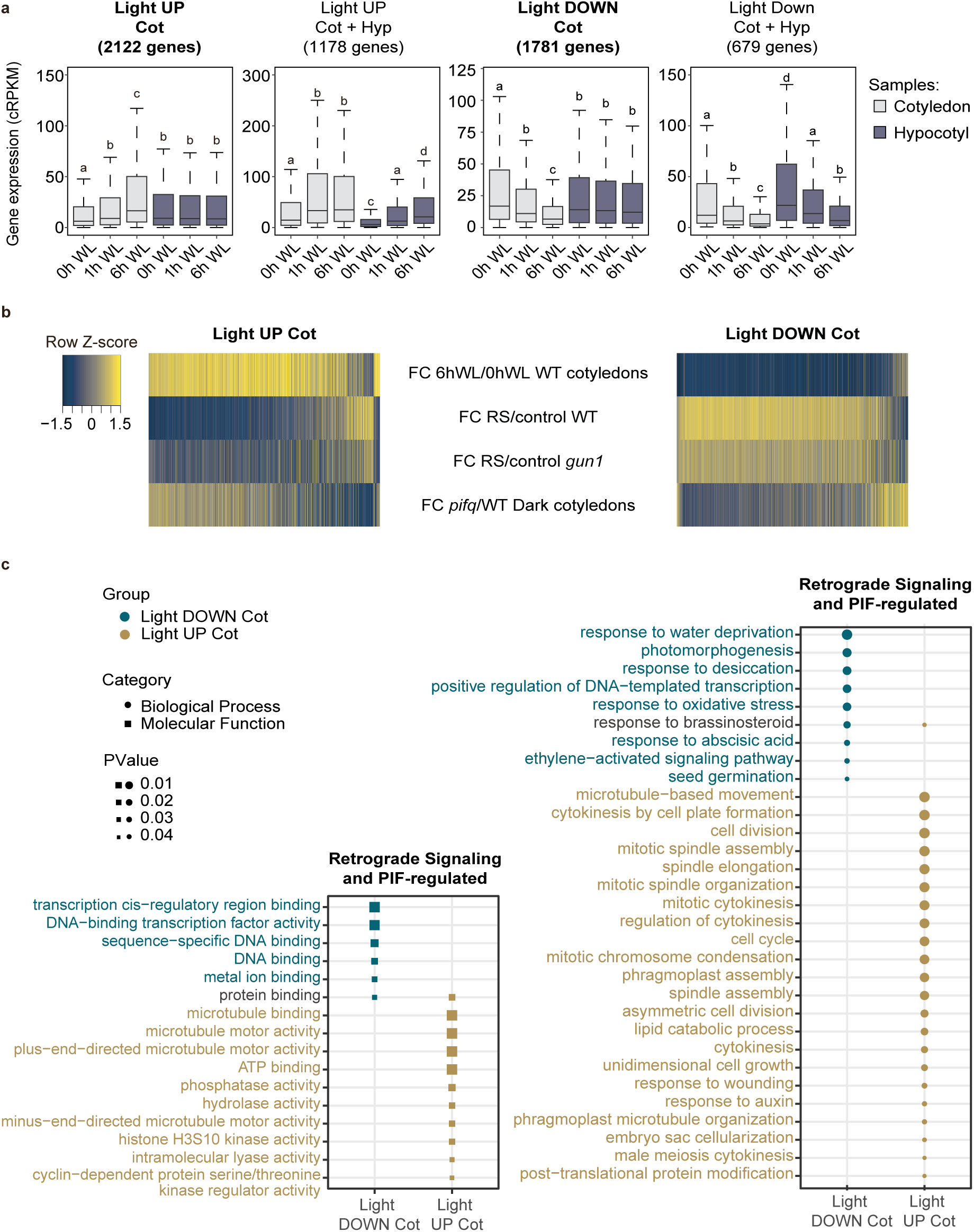
The cotyledon expression network is antagonistically regulated by phytochrome and retrograde signals with an enrichment in cell cycle genes. **(a)** Expression of genes in the “Light UP Cot”, “Light UP Cot+Hyp”, “Light DOWN Cot” and “Light DOWN Cot+Hyp” subsets in cotyledons or hypocotyls of dark-grown WT seedlings exposed to 1h or 6h of white light (WL). Letters denote statistically significant differences using one-way Kruskal-Wallis test (*P*<0.05). **(b)** Norflurazon antagonistically regulates the light-induced and PIF-repressed (left) and the light-repressed and PIF-induced (right) networks of “Light UP Cot” and “Light DOWN Cot” gene sets, in a partially GUN1-dependent manner. Heatmaps depicting fold-change (FC) expression of 2122 genes (left) and 1781 genes (right) in WT cotyledons after 6h of light exposure (top row), in WT and *gun1* after treatment with norflurazon (2 middle rows), and in cotyledons of dark-grown *pifq* compared to WT (bottom row). **(c)** Gene ontology terms of the “Light UP Cot” and “Light DOWN Cot” genes affected by at least two-fold both in *pifq* in the dark (Sun *et al*., 2016) and by norflurazon in the light (Zhao et al. 2019). About 10% of the “Light UP Cot” gene set (219 of 2122 genes) and 6.5% of the “Light DOWN Cot” (117 of 1781 genes) met both criteria. (a, c) Absolute and relative gene expression values were quantified from raw data submitted at the Sequence Read Archive under the accession numbers GSE79576 (Sun *et al*., 2016) and GSE110125 (Zhao *et al*., 2019).

Together, this analysis reveals that light activates a transcriptional program to reverse the PIFq-imposed gene expression landscape in dark-grown cotyledons, concomitant with the light-promoted switch in cotyledon growth axis and their expansion. Importantly, activation of GUN1-mediated RS antagonizes the effect of light, targeting both light induced and repressed gene networks, in agreement with the RS inhibition of cotyledon expansion in the light. We conclude that phytochrome and GUN1-mediated RS antagonistically regulate gene expression in cotyledons.

Next, to identify the most significant functional categories in each group, we first filtered the genes in each subset by selecting those that were affected by at least two-fold both in *pifq* in the dark (Sun et al. 2016) and by RS in the light (Zhao et al., 2019). About 10% of the “Light UP Cot” gene set (219 of 2122 genes) and 6.5% of the “Light DOWN Cot” (117 of 1781 genes) met both criteria (Supplemental Table 1). Gene Ontology analysis revealed that the “Light UP Cot” subset was highly enriched in cell cycle and cell division genes, whereas categories enriched in the “Light DOWN Cot” subset included transcription factor activity and hormone related genes among others (Figure 3c). The specificity of these categories in the cotyledon is underlined by the differential gene ontology profile of the “Light UP Cot+Hyp” and “Light DOWN Cot+Hyp” subsets (Supplemental Figure 1). Interestingly, these results suggest that cotyledon expansion might involve cell division, a long standing question in the field.

### Cotyledon epidermis growth during deetiolation relies solely on cell expansion

We next sought to elucidate the contribution of cell expansion and division within the light-induced expansion of the adaxial epidermis. In WT cotyledons grown in darkness, the epidermal pavement cells appeared elongated without lobes and aligned with the cotyledon long axis and shape (Figure 4a,b). After transferring to light, cells lost the preferential elongated growth and formed their characteristic lobes. After two days of red light, WT epidermis cells increased their size by 6.5-fold compared to 2-day-old dark cells (Figure 4a-d). Remarkably, epidermis pavement cells in dark-grown *pifq* seedlings were expanded and lobed, resembling cells in deetiolated WT cotyledons after 12 hours of red light (Figure 4a,b). In contrast, in the double mutant *phyAB*, pavement cells in the dark were similar to WT but were unable to expand in red light, maintaining the dark cell morphology of long cells without lobe formation (Figure 4c,d). We did detect some residual growth, likely due to the activity of phyC-E, in accordance to the lack of cotyledon expansion in seedlings deficient in all 5 phys (Hu *et al*., 2013).

**Figure 4.**
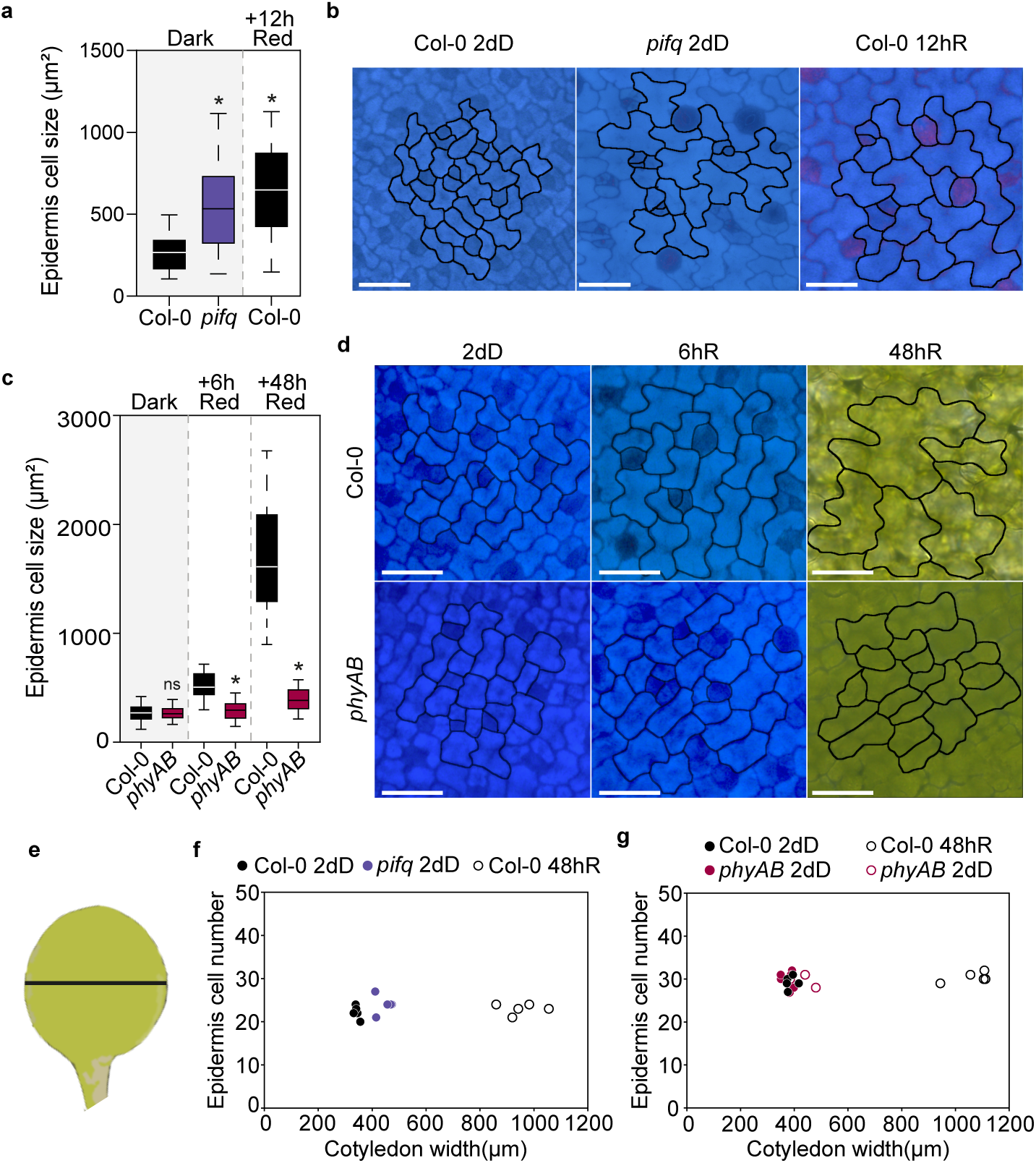
The PHY-PIF module regulates light-induced epidermis cell expansion during de-etiolation. **(a)** Quantification of epidermis cell size in WT and *pifq* in two-day-old seedlings (2dD) and WT seedlings transferred to red-light (R) for 12 hours. **(b)** Visual phenotypes of WT and *pifq* epidermis cells. **(c)** Quantification of epidermis cell size in WT and *phyAB* in two-day-old dark seedlings (2dD) and transferred to continuous red-light (R) for 6 and 48 hours. **(d)** Visual phenotypes of WT and *phyAB* epidermis cells in de-etiolation. Data in boxplots (a, c) indicate the first quartile, median, and third quartile of n≥125 cells pooled from at least 5 individual cotyledons. Whiskers indicate 5 to 95 percentile. Statistical differences relative to WT in each timepoint are indicated by an asterisk (Student t-test. *P*<0.05). ns, non-significant.**(e)** Representation of epidermis cell number relative to cotyledon width in WT and *pifq* (f), and in WT and *phyAB* (g) seedlings grown in the dark for two days (2dD) and in WT seedlings transferred to red light (R) for 48 hours. Each dot represents an individual cotyledon.

With regard to cell division, we only visually observed dividing cells in the relatively smaller subsets of cells surrounding stomata, probably byproducts of stomata formation as described (Houbaert *et al*., 2018; Smit *et al*., 2023). To provide quantitative evidence and confirm the absence of cell division in the rest of pavement cells, and given the major importance of width expansion as discussed above, we counted the number of cells present at the point of maximum width in the width axis of cotyledons, in a straight line from one edge of the cotyledon to the other, in 2dD seedlings and after 48h of exposure to Red light (48hR) (Figure 4 e-g). Next, for each measured cotyledon, we plotted the width in relation to the number of total cells in the width axis. Remarkably, despite the WT cotyledon width increase from ∼400 µm in the dark to almost 1200 µm after 48 hours of red light, the number of epidermal cells remains similar (Figure 4f). Dark-grown *pifq* also showed no difference in the number of cells compared to 2dD or 48hR WT (Figure 4f), despite of the wider and expanded *pifq* cotyledons in the dark (Figure 1). Similarly, we did not observe any changes in cell number in 2dD *phyAB* compared to 2dD WT, or to 48hR WT (Figure 4g). Together, these results suggest that cell expansion rather than division is the main contributor to epidermis development during de-etiolation, and that PIFs act as repressors of pavement cell expansion in the dark, a repression that is lifted upon exposure to red light through phy action.

### The palisade mesophyll experiences cell expansion and division during de-etiolation

Palisade cells, oriented with their long axis perpendicular to the cotyledon blade plane, can be visualized as circular structures in top-view microscopy in dark-and light-grown seedlings. In deetiolating WT, palisade cells rapidly expanded, from ∼150 in the dark to ∼240 um2 after 12h (Figure 5a,b), and to 600 um2 after 48h, an size increase of 4.5-fold (Figure 5c,d). In contrast, in dark-grown *pifq* palisade cells were clearly larger and displayed a size comparable to WT cells after 12 hours of red light (Figure 5a,b). In *phyAB*, palisade cell size in the dark was similar to WT, but the red-light induced expansion was strongly inhibited, showing only a slight growth after 48 hours (∼220 um2)(Figure 5c,d). These results resemble the expansion dynamics of the epidermis and the impact of the mutations in PIFq and phyA/B. However, in contrast to the epidermis, light induces palisade cell expansion without significantly changing the morphology (Figure 5b,d).

**Figure 5.**
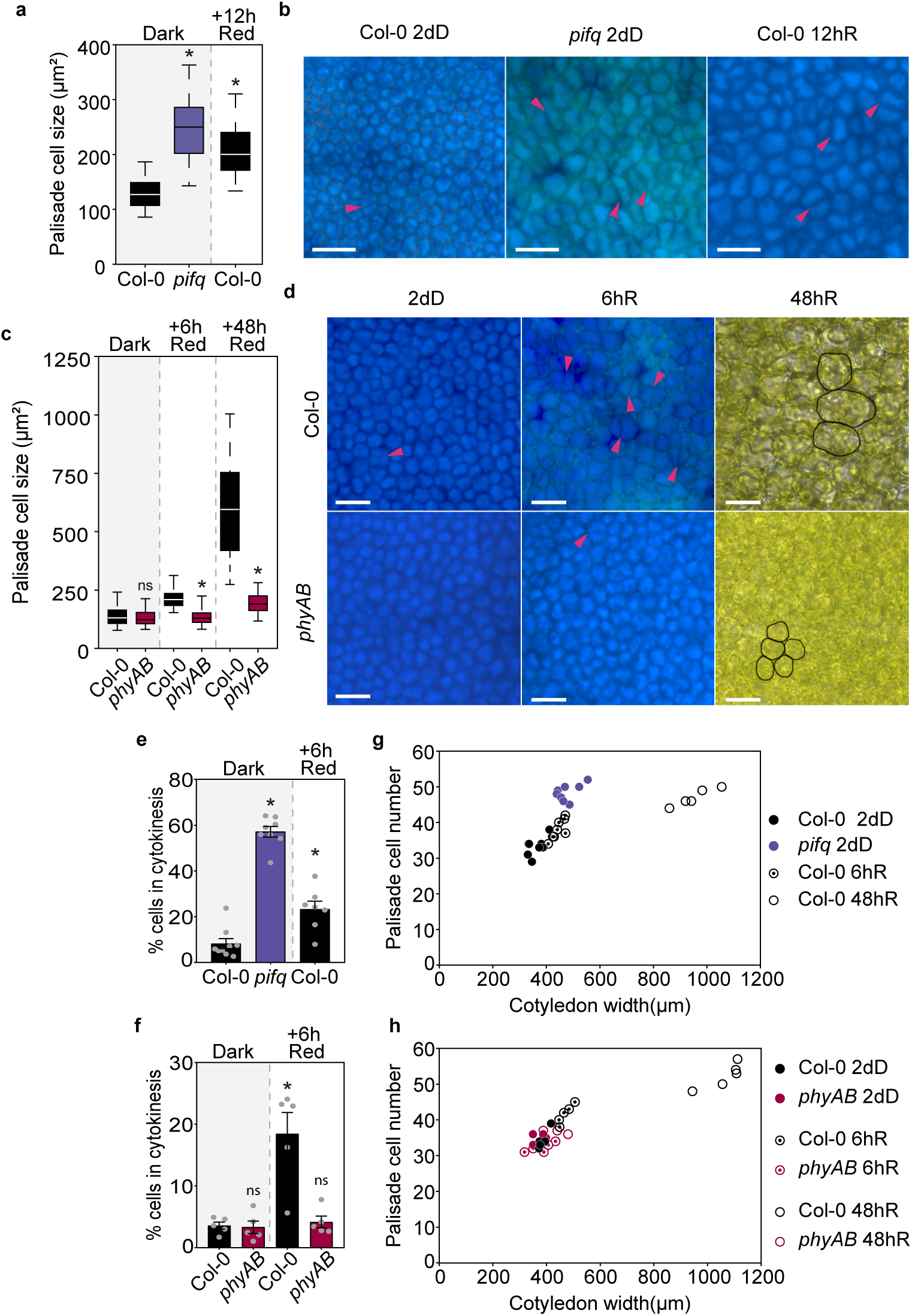
The PHY-PIF module regulates the light-induced palisade mesophyll cell growth during de-etiolation. **a)** Quantification of palisade cell size in WT and *pifq* in two-day-old seedlings (2dD) and WT seedlings transferred to red-light (R) for 12 hours. **(b)** Visual phenotypes of WT and *pifq* epidermis cells. **(c)** Quantification of palisade cell size in WT and *phyAB* in two-day-old dark seedlings (2dD) and transferred to continuous red-light (R) for 6 and 48 hours. **(d)** Visual phenotypes of WT and *phyAB* epidermis cells in de-etiolation. (a, c) Data in boxplots indicate the first quartile, median, and third quartile of n≥125 cells pooled from at least 5 individual cotyledons. Whiskers indicate 5 to 95 percentile. Statistical differences relative to WT in each timepoint are indicated by an asterisk (Student t-test. *P*<0.05). ns, non-significant. (b, d) Magenta arrows indicate visible cell division. **(e)** Percentage of palisade cells with a visible division plane in WT and *pifq* in two-day dark-grown seedlings (2dD) and WT seedlings transferred to red light (R) for 6 hours. **(f)** Representation of palisade cell number relative to cotyledon width in WT and *pifq* seedlings grown in the dark for two days (2dD) and WT seedlings transferred to red light (R) for 6 and 48 hours. **(g)** Percentage of palisade cells with a visible division plane in WT and *phyAB* in two-day dark-grown seedlings (2dD) and transferred to red light (R) for 6 hours. **(h)** Representation of palisade cell number relative to cotyledon width in WT and *phyAB* seedlings grown in the dark for two days (2dD) and transferred to red light (R) for 6 and 48 hours. Data in (e) and (g) indicate means ±SEM of at least 6 cotyledons. Asterisks indicate statistically significant differences relative to WT 2dD using Student t-test (*P*<0.05). In (e-h), each dot represents an individual cotyledon.

Interestingly, we visually observed dividing cells after the first hours of light. Indeed, a fraction of the cell population showed a clear cell wall division plane that divides the otherwise spherical palisade cell into two roughly identical cells (Figure 5b,d). Accordingly, the fact that deetiolating epidermis cells increase on average 6.5-fold in size compared to the 4.5-fold of palisade cells supports the notion that palisade cells must undergo division to support synchronized growth. To provide quantitative evidence, we assessed the percentage of visible palisade cells in cytokinesis in 2dD seedlings and upon red light exposure, in a centered square of 50×10^3^ µm^2^ that covered most of the cotyledon area. In WT seedlings, ∼5% of the palisade cells in the dark showed a visible division plane, that rapidly increased to ∼20% after 6 hours of red light (Figure 5e). Remarkably, in *pifq* the percentage of visible cells in cytokinesis in the dark was above 50% (Figure 5e), whereas in *phyAB* mutant seedlings the % of pavement cells was similar to WT in the dark and did not increase upon red light exposure (Figure 5f). Next, using the same approach as in the epidermis, we counted the number of cells in the width axis in different conditions and genotypes. Unlike the epidermis, the number of palisade cells in WT cotyledons increased from ∼35-40 cells in the dark, to ∼45 after 6h of red light, and ∼50-55 after 48 hours, an increase of ∼1.4X compared to dark (Figure 5g). Remarkably, in 2dD *pifq,* as a result of the increased cell division (Figure 5e), the number of total cells was similar to 2dD WT exposed to 48 hours of red light (Figure 5g). In the *phyAB* mutant, the lack of cotyledon expansion in red light was reflected in the number of palisade cells, which remained the same and similar to WT in the dark and after 48 hours of red light (Figure 5h). Together, these results indicate that during de-etiolation, cotyledon expansion involves division and expansion of palisade mesophyll cells, a process that is repressed by PIFs in the dark and activated upon exposure to red light through phy activity. Importantly, these results open the intriguing question of whether the increase in palisade cell division during deetiolation is induced by a PIF/phy-mediated light signal, and/or triggered by a mechanical stress signal due to the expansion forces of the adjacent cells, including the epidermal.

### Regulation of cotyledon expansion by cell-autonomous PIF and phyB activity in the epidermis

To start to shed light on the molecular mechanisms underlying cotyledon cell expansion and division, we took into consideration a recent report showing that PIF4 activity in epidermal cells affected cotyledon separation during skotomorphogenesis but not hypocotyl elongation (Kim *et al*., 2020). These data prompted us to hypothesize that epidermal PIFs could trigger epidermis expansion in the dark that might be enough to drive palisade cell division and growth, and overall cotyledon development. To test this we chose to focus on PIF1, as it has been described to be the major PIF contributor during skotomorphogenesis (Leivar *et al*., 2008; Pfeiffer *et al*., 2014). Transgenic plants expressing PIF1-GFP under the control of the epidermis-specific *MERISTEM LAYER1 (ML1)* promoter were generated in the *pifq* mutant background, and PIF1-GFP in these seedlings was detected in the epidermis pavement cells forming nuclear speckles, but not in the palisade (Figure 6a, Supplemental Figure 2) confirming tissue-specific accumulation. *PIF1* expression in the epidermis did not affect hypocotyl length and suppressed the cotyledon separation phenotype of *pifq* (Supplemental Figure 3) in accordance to the reported PIF4 data (Kim *et al*., 2020). Additionally, PIF1 activity in the epidermal cells was sufficient to suppress the expanded cotyledon phenotype of *pifq* in the dark, both in terms of area (Figure 6b-e) and also cell morphology, which displayed the elongated epidermal cells characteristic of etiolated WT cotyledons (Figure 6d). Moreover, no difference in epidermis cell number was observed when comparing *pifq* and *pML1::PIF1-GFP/pifq* (Figure 6f), in agreement with cotyledon expansion not requiring division in the epidermis. Strikingly, when examining the palisade, we observed that *pML1::PIF1-GFP/pifq* maintained elevated division rates comparable to *pifq* despite the unexpanded cotyledons (Figure 6g). In fact, both lines (*pifq and pML1::PIF1-GFP/pifq*) showed similar palisade cell number, causing *pML1::PIF1-GFP/pifq* to lose linearity in the correlation between cotyledon width and palisade cell number (Figure 6h).

**Figure 6.**
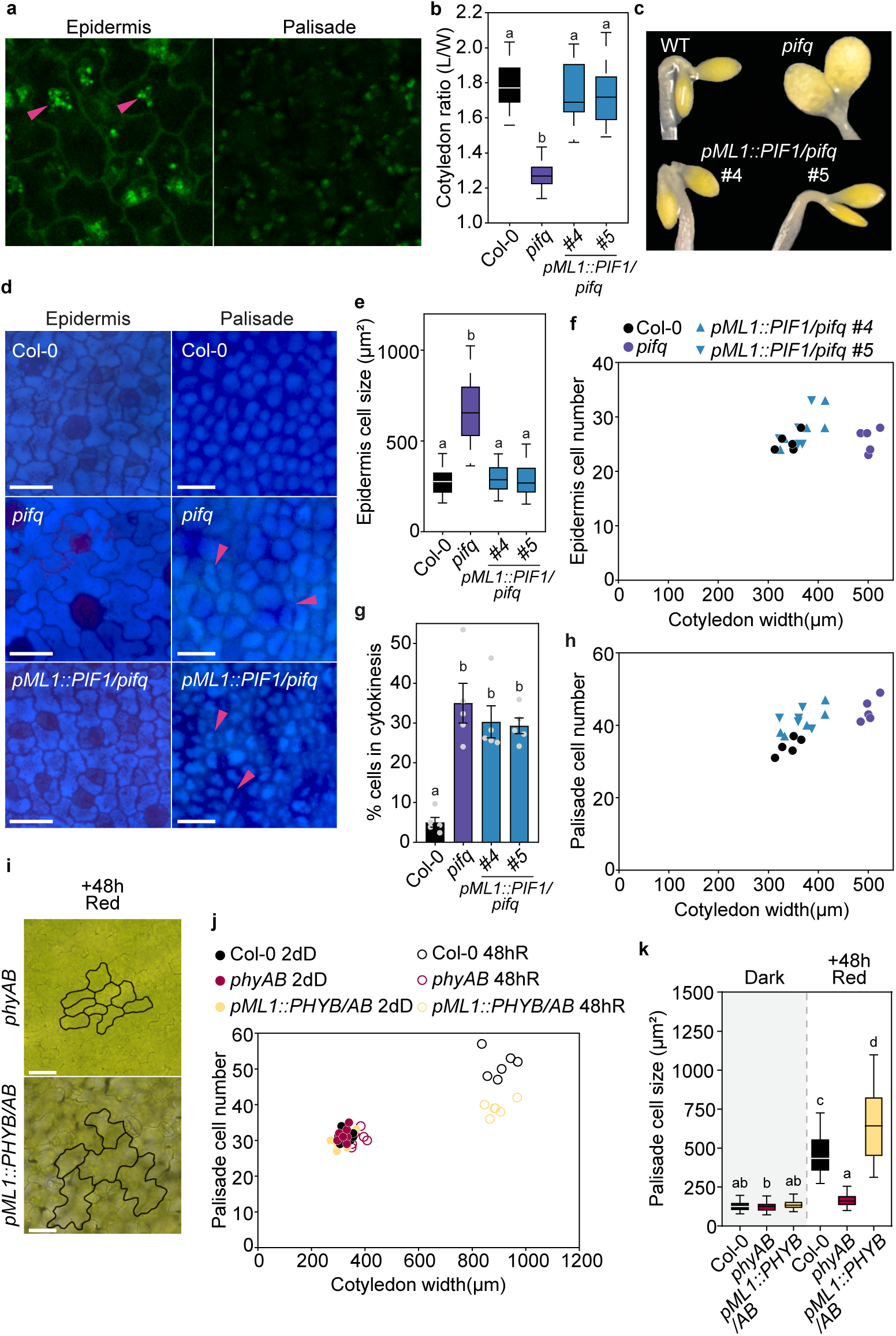
Epidermis-specific PIF1 expression reverts the *pifq* photomorphogenic cotyledon phenotype in the dark but not the elevated cell division in the palisade. **(a)** *PIF1-GFP* expression under the *pML1* promoter is detected by confocal microscopy in the nuclei of epidermis cells (magenta arrows) but not in the palisade. Seedlings were grown for 3d in dark and incubated for 16 hours in MG-132 (50µM). **(b)** Cotyledon ratio quantification of WT, *pifq*, and *pML1::PIF1-GFP* lines (*#4* and #*5*) in three-day-old dark-grown seedlings (3dD). Data are means±SEM of at least 25 seedlings. **(c)** Visual phenotypes of representative seedlings grown in (b). **(d)** Representation of epidermis cell number relative to cotyledon width in WT, *pifq*, and *pML1::PIF1-GFP #4/5* seedlings grown in the dark for two days. Each dot represents an individual cotyledon. **(e)** Epidermis cell size quantification in WT, *pifq*, and *pML1::PIF1-GFP #4/5* seedlings grown in the dark for two days. Boxplots indicate the first quartile, median, and third quartile. Whiskers indicate 5 to 95 percentiles. **(f)** Representation of palisade cell number relative to cotyledon width in WT, *pifq*, and *pML1::PIF1-GFP #4/5* seedlings grown in the dark for two days. Each dot represents an individual cotyledon. **(g)** Percentage of palisade cells in visible division in WT, *pifq*, and *pML1::PIF1-GFP #4/5* cotyledons grown in the dark for two days. Data are means ±SEM of at least 6 cotyledons. **(h)** Visual phenotypes of epidermis and palisade cells in WT, *pifq*, and *pML1::PIF1-GFP #4/5* cotyledons grown in the dark for two days. (b,e,g) Letters denote the statistically significant differences using one-way ANOVA followed by post hoc Tukey’s test (*P*<0.05). (i) Visual phenotypes of epidermis cells in *phyAB* and *pML1::PHYB/AB* cotyledons grown in the dark for two days and transferred to red light for 48h hours. (j) Representation of palisade cell number relative to width in WT, *phyAB* and *pML1::PHYB/AB* seedlings grown in the dark for two days (2dD) and transferred to red light (R) for 48 hours. Each dot represents an individual cotyledon. (k) Palisade cell size quantification in WT, *phyAB* and *pML1::PHYB/AB* seedlings grown in the dark for two days (2dD) and transferred to red light (R) for 48 hours. Data in boxplots indicate the first quartile, median, and third quartile of n≥125 cells pooled from at least 5 individual cotyledons. Whiskers indicate 5 to 95 percentile. Letters denote statistically significant differences using two-way ANOVA followed by post hoc Tukey’s test (*P*<0.05).

We also attempted to generate transgenic plants to express *PIF1* under a mesophyll-specific promoter, an objective that recently proved to be challenging (Kim *et al*., 2020). We discarded the commonly used *CHLOROPHYLL A/B BINDING PROTEIN 3* (*CAB3*) or *IQ-DOMAIN-22* (*pIQD22*) promoters as both have been recently shown to also drive epidermal activity in cotyledons, probably through leaky expression (Kim *et al*., 2020; Procko *et al*., 2022), and expressed *PIF1:mScarlet* under the control of the *SQALENE MONOOXYGENASE 6* promoter (*pSQE6*), shown to drive expression in the palisade and the lowermost spongy layer in leaves (Procko *et al*., 2022). However, we were not able to detect *PIF1* expression in our transformed seedlings, probably due to very low expression levels in the cotyledon as previously reported (Sun *et al*., 2016).

To further understand the epidermis/palisade interplay in the regulation of cotyledon expansion, we next examined available *pML1::PHYB-GFP* expressing lines in the *phyAphyB* background (*pML1::PHYB-GFP/AB)* (Kim *et al*., 2016). *phyB* expression in the epidermis was sufficient to drive cotyledon expansion in response to light (Figure 6i, j), showing the characteristic lobe morphology (Figure 6i). Strikingly, when examining the palisade, we observed that *pML1::PHYB-GFP/AB* palisade cells were abnormally larger (Figure 6k), possibly due to the mechanical stress exerted by the growing epidermis, and exhibited highly reduced cell division (Figure 6j).

Together, these results underscore the importance of the epidermis in cotyledon growth, suggesting that a phy/PIF cell-autonomous activity in the epidermis can govern the extent of cotyledon expansion. Importantly, our results also highlight that the regulation of the palisade cell division does not need a light signal from the epidermis, although mechanical stress from an expanding epidermis can trigger some palisade cell division and both signals probably co-exist. Our results strongly suggests that the light-induced palisade division rate is cell-layer autonomous, and reveal a cell-autonomous PIF activity in the palisade to repress cell division in the dark.

## DISCUSSION

We have shown here that light-induced cotyledon expansion involves a switch in the growth direction that is antagonistically regulated by light through the PIF/phy system and by GUN1-mediated retrograde signaling. Our work examines the contribution of cell expansion and division in the epidermis and the mesophyll, and shows for the first time that cotyledon palisade expansion involves cell division. This aligns with enrichment of cell division-related genes antagonistically regulated by the PIF/phy system and RS specifically in the cotyledon. Furthermore, using mutant lines specifically expressing PIF1 and phyB in the epidermis, our results show that epidermis expansion can drive cell growth in the palisade, whereas mesophyll cell division is mostly regulated by light at tissue-specific level. These results, together with previous data, support a model whereby cotyledon expansion during deetiolation relies on a PIF/phy-regulated switch in growth direction from longitudinal in the dark to transversal in the light, and involves distinct growth strategies for the epidermis (based on cell expansion) and mesophyll layers (based on cell expansion and division) (Figure 7). Light-induced cotyledon growth requires and coincides temporarily with etioplast-to-chloroplast development, which ensures availability of the maximum light-capturing surface for photosynthesis when the light intensity is adequate. The system is exquisitely sensitive to the prevailing light environment, and potentially damaging high light conditions activate GUN1-mediated RS to inhibit cotyledon expansion and reduce the exposed area susceptible to damage (Figure 7). Hence, RS acts as a developmental brake that could be critical under conditions such as fluctuating light intensity under a canopy with locally high irradiance sunflecks (Durand *et al*., 2021), or as a function of cloud coverage (Schneider et al., 2019).

**Figure 7.**
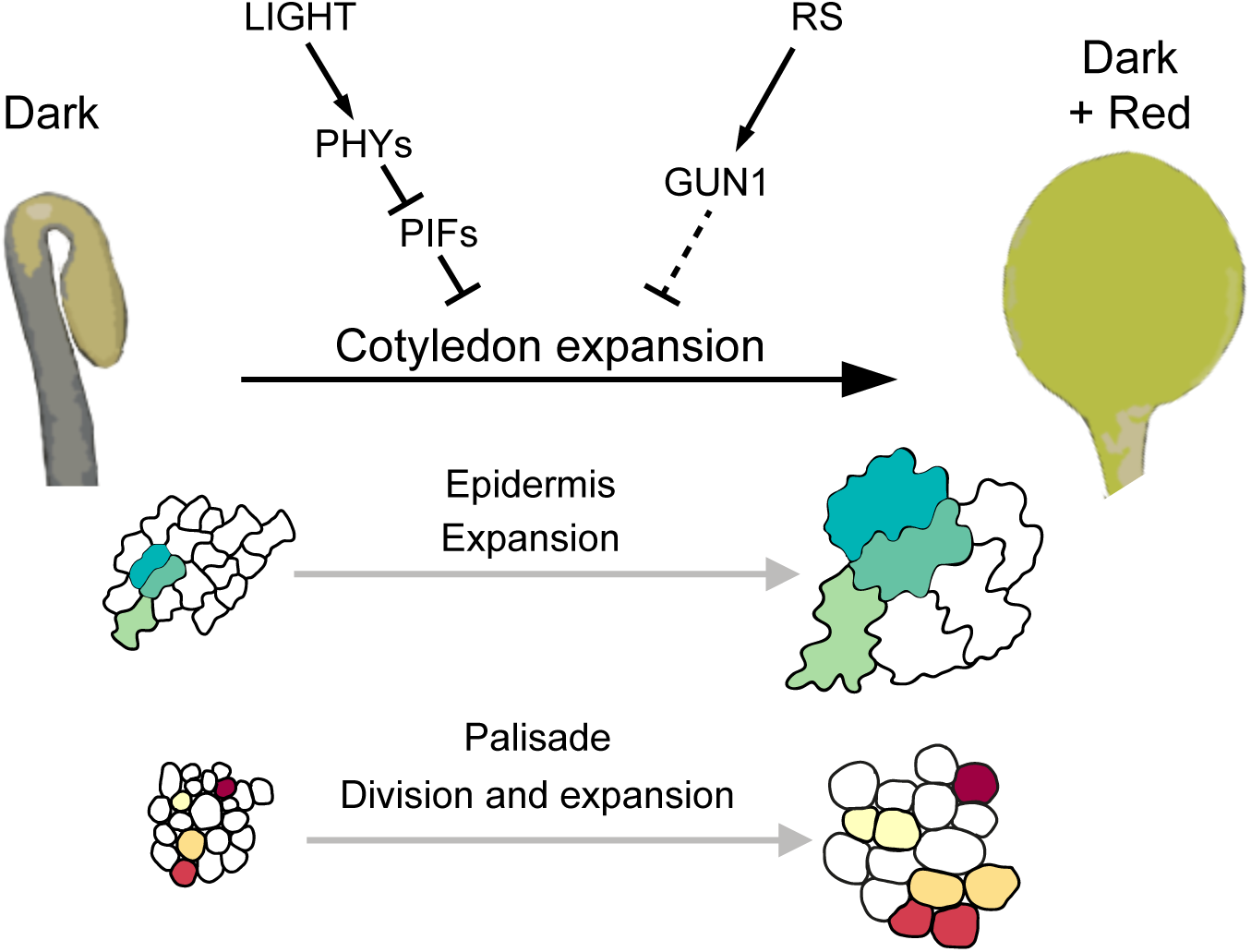
Model of PIF- and GUN1-mediated regulation of cotyledon expansion during seedling deetiolation. Schematic model depicting PIF-mediated repression of cotyledon expansion in the dark. Upon exposure to light, active phys repress PIF activity and trigger PIF degradation, and cotyledons start to rapidly expand. Cotyledon growth involves epidermis cell expansion and palisade cell division and expansion. Cotyledon growth takes place coinciding with etioplast-to-chloroplast development. If chloroplast biogenesis is disrupted, GUN1-mediated retrograde signaling inhibits cotyledon expansion.

We have established a novel framework to study how cotyledon expansion is regulated during seedling deetiolation. Cotyledon expansion has been traditionally overlooked in light signaling and seedling de-etiolation research, where hypocotyl elongation and cotyledon separation to a lesser extent have been extensively and preferentially used as phenotypic readouts. The advantages of hypocotyl as a simple model thanks to its unidirectional growth based on cell expansion (Gendreau *et al*., 1997) and the similar morphology of its cell types, has contributed greatly to the advancement of our knowledge in light signaling (Zhao *et al*., 2020), de-etiolation (Jedynak *et al*., 2022), light-hormonal interplay (Zhang *et al*., 2015), circadian clock (Martín *et al*., 2020), and shade avoidance (Sharma *et al*., 2023) among others. However, the complexity of seedling development has long been evident in the opposite effect of light on hypocotyl and cotyledon development, where light inhibits hypocotyl elongation while promoting expansion of cotyledons (Bou-Torrent et al., 2008). This opposite regulation is crucial for seedling survival, and a detail understanding of cotyledon development can provide insights into the underlaying mechanisms. We have shown that light-induced cotyledon expansion relies on a switch of organ growth direction, from longitudinal to transversal (Figure 1). Because plant cells are enclosed by a rigid cell wall and cannot migrate, the control of directional cell growth determines morphogenesis and organ geometry. This establishment and coordination of plant growth and development relies on both mechanical as well as biochemical signals governing cell growth (Jonsson et al., 2022). The regulation of growth direction remains relatively unclear, but it is in part controlled by the orientation of the cortical microtubules (Kirchhelle *et al*., 2019). Interestingly, some of the enriched gene-ontology terms found in our transcriptomic analysis of the light/PIF-induced and RS-regulated genes in the cotyledon (Figure 3) are related to microtubule movement and organization. A recent report has shown that mutants in the OVATE FAMILY PROTEIN (OFP) of transcriptional repressors exhibit long cotyledons when grown in the light due to enhanced cell elongation (X. Zhang *et al*., 2020). OFP proteins regulate cell growth direction in cotyledons by interacting with TONNEAU2 (TON2) to modulate microtubule orientation. TON2 is able to interact with LONGIFOLIA (LNG) proteins (Drevensek *et al*., 2012), which positively regulate polar cell elongation in the leaf length axis to determine morphology, and have been shown to be directly regulated by PIFs (Hwang *et al*., 2017). Interestingly, the OFP-TON2 interaction is mediated by brassinosteroid levels (X. Zhang *et al*., 2020), and LNGs promote growth by activating the auxin pathway (Hwang *et al*., 2017). Growth directionality in several organs is determined by hormone gradients. In apical hooks, the PIN3 and PIN4 auxin carriers drive auxins toward the outermost convex side of the hook, draining auxin concentration in the inner concave side and promoting hook opening (Žádníkova *et al*., 2010; Abbas *et al*., 2013). In roots, auxin gradients control root growth, cytokinin acts antagonistically to regulate cell division, and gibberellins delimit the front between cell expansion and cell division (Beemster and Baskin, 2000; Grieneisen *et al*., 2007; Achard *et al*., 2009). Additionally, cytokinin gradients generate asymmetric cell division to generate root bending in hydrotropic responses (Chang *et al*., 2019). In leaves, WOX proteins (WUSCHEL-RELATED HOMEOBOX) regulate auxin biosynthesis in leaf margins, creating an auxin gradient that promotes leaf width-specific growth (Z. Zhang *et al*., 2020). In all, the connection between organ growth directionality, cell elongation, microtubule orientation, and hormone pathways and/or gradients has been extensively studied in the regulation of leaf shape, hypocotyl elongation, apical hook development or root growth (Jonsson et al., 2022). However, whether cotyledon expansion relies on a similar regulatory coordination of mechanical and hormone signals and gradients is unknown. Our finding that hormone categories (such as brassinosteroid, ABA and ethylene) are enriched in our transcriptomic analysis of the light/PIF- and RS-regulated genes during cotyledon expansion together with microtubule terms (Figure 3c) suggests a similar general regulatory framework. Interestingly, all three hormones have been previously shown to affect microtubule orientation (Hamant and Traas, 2010). In addition, organ-specific transcriptomic analysis identified the auxin-regulated *Small Auxin Up RNA* (*SAUR*) genes whose transcripts are light-induced in cotyledons and/or repressed in hypocotyls (Sun *et al*., 2016). Importantly, PIFs were shown to directly bind these *SAURs* and differentially regulate their expression in cotyledons and hypocotyls possibly through interaction with organ-specific cofactors (Sun *et al*., 2016), suggesting overall that auxin might play a role in the regulation of cotyledon expansion. Whether auxin and/or SAURS might affect microtubule orientation is currently unknown. Future studies to explore the mechanical and biochemical signaling interplay in cotyledons will be of great interest, as well as to determine whether there is a tissue-level control differentiating cell growth direction of epidermis and palisade cells, for which the cotyledon provides an excellent study system.

The effect of light during seedling deetiolation on the shape of epidermal and palisade cells is clearly distinct. Epidermal cells acquire lobed puzzle-shape forms as they expand (Figure 4). This complex shape has been proposed to reduce mechanical stress in the cell wall during isotropic growth and prevent bulging due to the turgor pressure (Sapala *et al*., 2018). In contrast, cells in the palisade layer expand keeping the original rectangular shape (Figure 5). Interestingly, our finding that palisade cells undergo cell division during cotyledon expansion might also represent a strategy to minimize and cope with the mechanical stress. Indeed, the orientation of cell division has been proposed to align with the direction of maximal mechanical stress to produce smaller cells that will have a reinforced cell wall in the orientation of maximum tension (Robinson, 2021). In addition, although the regulation of chloroplast division and the integration with cell division and expansion is still poorly understood (Jarvis and López-Juez, 2013), we propose that cell division in the palisade may indirectly allow an increase in the number of chloroplasts in the mesophyll. This would contribute to increase the photosynthetic capacity of the expanding cotyledon during seedling deetiolation, although to our knowledge no studies have addressed this directly.

Our results showing that the light regulation of palisade cell division seems to be largely independent of the epidermis, opens the question of how these two layers coordinate growth during cotyledon expansion. In our experiments expressing PIF1 and phyB under the control of the epidermis-specific promoter pML1, epidermis and palisade extension are coordinated (Figure 6). It seems therefore that the mechanical force exerted by the epidermis works as a coordinating signal for the palisade, as suggested in previous work (Verger *et al*., 2018). However, our finding that cell division in the palisade largely escapes this coordination suggests that the regulation of cell expansion and division are likely uncoupled, being cell division more strictly regulated by light at tissue-specific level.

Finally, our work highlights the use of cotyledon development in response to light as a new model system to study organ shape regulation, a fundamental challenge in biology. Cotyledons provide a powerful model to address unanswered questions for which other organs like hypocotyls are not suitable, such as the coordination of different cell layers during growth, light-induced multidimensional expansion, or the coordination of the individual axes in growth. Moreover, although cotyledon expansion and leaf growth present considerable differences (for example the presence of a leaf primordia that initially grow by cell division and is progressively substituted by cell expansion), leaf and cotyledon cell layer structures are equivalent, and cotyledons can help addressed open questions in leaf development such as the interconnection between division and expansion or the interplay between the different cell layers (Gonzalez *et al*., 2012), using the tools and approaches described here.

## EXPERIMENTAL PROCEDURES

### Plant material

*Arabidopsis thaliana* seeds used in this manuscript include the previously described Col-0, *phyA-211*(Reed et al., 1994), *phyB-9* (Reed *et al*., 1994), *phyA-211phyB-9* (Reed *et al*., 1994), *pifq* (Leivar *et al*., 2008), and *pML1::phyB-GFP/phyAphyB* (*pML1::PHYB-GFP/AB* (Kim *et al*., 2016). To generate the *pML1::PIF1-GFP* transgenic lines, the *MERISTEM LAYER-1* promoter described in (van Es *et al*., 2018) was amplified using primers indicated in Table 1 (EMP1730+EMP1731) and cloned into pDONR-P4-P1R using the BP recombination reaction (Gateway, Invitrogen). Similarly, *PIF1* coding sequence was amplified (using primers EMP1732+EMP1732, see Supplemental Table 2.) and cloned into pDONR221 and the GFP coding sequence into pDONR-P2R-P3. LR recombination reaction (Gateway ®, Invitrogen) was performed to generate in-frame *pML1::PIF1-GFP* constructs into the pH7m34GW expression vector that was transformed into the *pifq* background.

### Growth conditions

Seeds were surface sterilized with 20% bleach and 0.5% sodium dodecyl sulfate (SDS) for 10 minutes and rinsed five times with distilled sterile water. Seeds were plated on half-strength Murashige and Skoog (0.5xMS) medium without sucrose, stratified at 4°C in the dark for 4d, and exposed to white light (100 µE) at 21°C for 3h to induce germination. Seedlings were then kept in the dark at 21°C for 2d and transferred to continuous red light (20 µE) at 21°C for the specified hours in each experiment. For dark-extended experiments, seedlings were kept in the dark at 21°C for 2, 3, 4, or 5 days. For lincomycin treatments, the medium was supplemented with 0.5mM lincomycin (Sigma L6004) (Sullivan and Gray, 1999). Red light were provided by PHILIPS GreenPower research module in deep red (maximum at 660 nm). Light intensity was measured with a built-in LI-COR LI-190R Quantum Sensor in the red growth chambers (Aralab) and with a hand-held SpectraPen mini (Photon Systems Instruments) for the white light growth chamber (Aralab).

### Cotyledon imaging measurements

To quantify cotyledon expansion, seedlings were gently pushed against the media, and cotyledons were gently separated using tweezers and photographed using a digital camera (Nikon D7000). Measures were performed using NIH image software (Image J, National Institutes of Health). The length (from the cotyledon base at the petiole attachment point to the cotyledon tip in a straight line) and the width (from one lateral border to the other in a straight line in the wider point of the cotyledon) were measured. Only one cotyledon per seedling was measured. The cotyledon ratio was defined by dividing length/width.

### Cell imaging and measurements

Cell imaging was performed using the DM6 epifluorescence microscope (Leica). Cotyledons were cut and mounted in water. Multiple images from each cotyledon were obtained using a 63X lens and later merged to generate composite images of the whole cotyledon using the LASX software (Leica). For dark, 6, and 12 hours of red light exposure, cotyledon images were captured using the UV excitation range A-filter cube (340-380nm) and a suppression filter LP425, providing high contrast to the cell walls without the need for chemical staining. For 6 and 12h red light images, the “red channel” from the images was deactivated to eliminate the chlorophyll signal. In 24 and 48-hour red light samples, images were captured using bright field imaging, as high chlorophyll content interferes with the signal obtained with the A-filter cube.

For cell size (epidermis and palisade) measurements, at least 25 cells per cotyledon in at least 5 cotyledons (n≥125) were measured from the central part of the cotyledon. In the palisade, only non-dividing cells were measured. Measures were performed using NIH image software (Image J, National Institutes of Health). For cell counting, a straight line following the width axis in the middle of the cotyledon was drawn, and all cells crossed by the line were counted. For the quantification of the percentage of palisade cells in cytokinesis, the total number of cells and those in visible cytokinesis were counted in an area of 50 x10^3^ µm^2^ in the center of the cotyledon.

### Fluorescence microscopy

For PIF1-GFP epidermal detection, *pML1::PIF1-GFP* seedlings were grown for 3d in the dark. Then, seedlings were incubated in the dark for 16 hours in liquid MS media with MG-132 (50µM) (Calbiochem) protease inhibitor. Lines were visualized using the SP5 confocal microscope (Leica) with an excitation argon laser (488nm) and emission range (500-600nm).

### Statistical analysis

Differences in cotyledon ratio, percentage of cells, and cell size between two genotypes were analyzed by Student t-test, and significantly different pairs (*P*<0.05) were represented by asterisks. To identify differences in cotyledon ratio, cell size, percentage of cells or relative gene expression between multiple genotypes in one light conditions, data were analyzed using one-way ANOVA followed by post hoc Tukey’s test (*P*<0.05), and results were represented by letters. To identify differences in cotyledon ratio or cell size between multiple genotypes and light conditions, data were analyzed using two-way ANOVA followed by post hoc Tukey’s test (*P*<0.05), and results were represented by letters.

### Gene Expression Analyses

To detect *PIF1* levels in the generated *pML1::PIF1-GFP* lines, qRT-PCR was performed as described previously (Veciana *et al*., 2022). Briefly, seedlings were grown in the dark and total RNA was extracted using Mawxell RSC plant RNA Kit (Promega). One µg of total RNA was treated with DNase I (Promega) according to the manufacturer’s instructions. First-strand cDNA synthesis was performed using the NZYtech First-strand cDNA Synthesis Kit (NZYtech), and 2 µl of 1:25 diluted cDNA with water was used for real-time PCR (LightCycler 480 Roche) using SYBR Premix Ex Taq (Takara) and primers at 300nM concentration. Gene expression was measured in three independent biological replicates, and at least two technical replicates were done for each of the biological replicates. *PP2A* (AT1G13320) was used for normalization (Shin et al., 2007). Primer sequences are described in Supplementary Table 2.

### RNAseq quantification of gene expression levels and Gene Ontology analyses

Quantification of total mRNA levels from public sequencing data (GSE79576; Sun et al. 2016 and GSE110125; Zhao et al., 2019) used in Figure 3 and Supplemental Figure 1 was performed with *vast-tools* v2.5.1 (Martín *et al*., 2021). This tool provides the corrected-for-mappability RPKMs (cRPKMs), corresponding to the number of mapped reads per million mapped reads, divided by the number of uniquely mappable positions of the transcript (Labbé *et al*., 2012). The list of differentially expressed genes in each condition: WT Light, *pifq* Dark and Norflurazon-treated seedlings were previously reported (Sun *et al*., 2016 and Zhao *et al*., 2019). Significantly enriched biological processes and molecular functions among the different sets of genes were defined using the functional annotation classification system DAVID (Huang *et al*., 2007).

## ACCESSION NUMBERS

Raw sequencing data used in this study were obtained from the Sequence Read Archive (accession numbers GSE79576 and GSE110125).

## ACKNOWLEDGEMENTS

We are grateful to Giltsu Choi for sharing seeds of the *pML1:PHYB/AB* line. This work was supported by grants from FEDER/Ministerio de Ciencia, Innovación y Universidades – Agencia Estatal de Investigación (Project References BIO2015-68460-P, PGC2018-099987-B-I00, and PID2021-122288NB-I00 to E.M.; and RYC2020-030160-I and PID2021-125223NA-I00 to G.M.), and from the CERCA Programme/Generalitat de Catalunya (Project References 2017SGR-718 and 2021SGR-792 to E.M.; and 2021SGR-873 to G.M.). We acknowledge financial support from the Spanish Ministry of Economy and Competitiveness, through the ‘Severo Ochoa Programme for Centres of Excellence in R&D’ 2016–2019 (SEV-2015-0533) and CEX2019-000902-S funded by MCIN/AEI/10.13039/501100011033.

## SHORT LEGENDS FOR SUPPORTING INFORMATION

**Supplemental Table 1.** Gene lists corresponding to the light-regulated gene sets defined in Figure 3. These genes were identified based on their light-responsiveness in cotyledon and hypocotyl organs and their regulation by PIFs (Sun *et al*., 2016) and retrograde signaling (RS; Zhao *et al*., 2019).

**Supplemental Table 2.** Primer sequences used for cloning and gene expression analysis. **Supplemental Figure 1.** Gene Ontology (GO) enrichment of the “Light UP Cot+Hyp” and “Light DOWN Cot+Hyp” subsets regarding biological process and molecular function categories.

**Supplemental Figure 2.** Epidermis-specific *PIF1-GFP* Characterization of independent *pML1::PIF1-GFP/pifq* lines.

**Supplemental Figure 3.** Epidermis-specific *PIF1-GFP* expression complements the constitutively photomorphogenic cotyledon phenotype of *pifq* in *pML1::PIF1-GFP/pifq* lines but not the short hypocotyl.

## TABLES

**Supplemental Table 1.** Gene lists corresponding to the light-regulated gene sets defined in Figure 3. These genes were identified based on their light-responsiveness in cotyledon and hypocotyl organs and their regulation by PIFs (Sun et al., 2016) and retrograde signaling (RS; Zhao et al., 2019).

**Supplemental Table 2.** Primer sequences used for cloning and gene expression analysis.

## FIGURE LEGENDS

**Supplemental Figure 1.** Gene Ontology (GO) enrichment of the “Light UP Cot+Hyp” and “Light DOWN Cot+Hyp” subsets regarding biological process and molecular function categories.

**Supplemental Figure 2. Epidermis-specific *PIF1-GFP* Characterization of independent *pML1::PIF1-GFP/pifq* lines. (a)** Quantification of cotyledon ratio (length/width) in WT, *pifq*, and independent *pML1::PIF1-GFP/pifq* transgenic lines. Data in boxplots indicate the first quartile, median, and third quartile of n≥20 seedlings. Whiskers indicate 5 to 95 percentile. Letters denote the statistically significant differences using one-way ANOVA followed by post hoc Tukey’s test (*P*<0.05). **(b)** *PIF1* expression relative to Col-0 set at 1 in 2dD Col-0, *pifq*, and independent *pML1::PIF1-GFP/pifq* transgenic lines. Data are the means ± SE of biological triplicates (n = 3). **(c)** Detection of *PIF1-GFP* accumulation by confocal microscopy in the nuclei of epidermis cells (magenta arrows). Seedlings were grown for 3 days in the dark and incubated for 16 hours in MG-132 (50µM).

**Supplemental Figure 3. Epidermis-specific *PIF1-GFP* expression complements the constitutively photomorphogenic cotyledon phenotype of *pifq* in *pML1::PIF1-GFP/pifq* lines but not the short hypocotyl. (a)** Expression of epidermis-specific *PIF1-GFP* does not rescue the short hypocotyl phenotype of *pifq*. Hypocotyl length of 3dD WT, *pifq*, and two independent *pML1::PIF1-GFP/pifq* lines. Data in boxplots indicate the first quartile, median, and third quartile of n≥20 seedlings. Whiskers indicate 5 to 95 percentile. Letters denote the statistically significant differences using one-way ANOVA followed by post hoc Tukey’s test (*P*<0.05). **(b)** Visual phenotypes of representative seedlings grown as in (a), showing complementation of the cotyledon expansion and separation phenotype.

## REFERENCES

1. Abbas, M., Alabadí, D. and Blázquez, M.A. (2013) ‘Differential growth at the apical hook: All roads lead to auxin’, Frontiers in Plant Science, 4(NOV), p. 64424. Available at: 10.3389/fpls.2013.00441.

2. Achard, P., et al. (2009) ‘Gibberellin signaling controls cell proliferation rate in Arabidopsis’, Current biology : CB, 19(14), pp. 1188–1193. Available at: 10.1016/j.cub.2009.05.059.

3. Bean, G.J., et al. (2002) ‘Tissue patterning of Arabidopsis cotyledons’, New Phytologist, 153(3), pp. 461–467. Available at: 10.1046/J.0028-646X.2001.00342.X.

4. Beemster, G.T.S. and Baskin, T.I. (2000) ‘Stunted plant 1 mediates effects of cytokinin, but not of auxin, on cell division and expansion in the root of Arabidopsis’, Plant physiology, 124(4), pp. 1718–1727. Available at: 10.1104/PP.124.4.1718.

5. Borsuk, A.M., et al. (2022) ‘Structural organization of the spongy mesophyll’, The New Phytologist, 234(3), p. 946. Available at: 10.1111/nph.17971.

6. Bou-Torrent, J., Roig-Villanova, I. and Martínez-García, J.F. (2008) ‘Light signaling: back to space’, Trends in Plant Science, 13(3), pp. 108–114. Available at: 10.1016/j.tplants.2007.12.003.

7. Cantón, F.R. and Quail, P.H. (1999) ‘Both phyA and phyB Mediate Light-Imposed Repression of PHYA Gene Expression in Arabidopsis’, Plant Physiology, 121(4), p. 1207. Available at: 10.1104/pp.121.4.1207.

8. Carter, R., et al. (2017) ‘Pavement cells and the topology puzzle’, Development (Cambridge), 144(23), pp. 4386–4397. Available at: 10.1242/dev.157073

9. Chang, J., et al. (2019) ‘Asymmetric distribution of cytokinins determines root hydrotropism in Arabidopsis thaliana’, Cell Research 2019 29:12, 29(12), pp. 984–993. Available at: 10.1038/s41422-019-0239-3.

10. Chen, C., et al. (2012) ‘Light-Regulated Stomatal Aperture in Arabidopsis’. Available at: 10.1093/mp/sss039.

11. Debrieux, D. and Fankhauser, C. (2010) ‘Light-induced degradation of phyA is promoted by transfer of the photoreceptor into the nucleus’, Plant molecular biology, 73(6), pp. 687–695. Available at: 10.1007/S11103-010-9649-9.

12. Drevensek, S., et al. (2012) ‘The Arabidopsis TRM1–TON1 Interaction Reveals a Recruitment Network Common to Plant Cortical Microtubule Arrays and Eukaryotic Centrosomes’, The Plant Cell, 24(1), p. 178. Available at: 10.1105/TPC.111.089748.

13. Durand, M., et al. (2021) ‘Sunfleck properties from time series of fluctuating light’, Agricultural and Forest Meteorology, 308–309, p. 108554. Available at: 10.1016/J.AGRFORMET.2021.108554.

14. van Es, S.W. et al. (2018) ‘Novel functions of the Arabidopsis transcription factor TCP5 in petal development and ethylene biosynthesis’, The Plant Journal, 94(5), p. 867. Available at: 10.1111/TPJ.13904.

15. Gendreau, E., et al. (1997) ‘Cellular Basis of Hypocotyl Growth in Arabidopsis thaliana’, Plant Physiology, 114(1), pp. 295–305. Available at: 10.1104/PP.114.1.295.

16. Gommers, C.M.M. and Monte, E. (2018) ‘Seedling Establishment: A Dimmer Switch-Regulated Process between Dark and Light Signaling’, Plant Physiology, 176(2), pp. 1061–1074. Available at: 10.1104/pp.17.01460.

17. Gonzalez, N., Vanhaeren, H. and Inzé, D. (2012) ‘Leaf size control: complex coordination of cell division and expansion.’, Trends in Plant Science, 17(6), pp. 332–340. Available at: 10.1016/J.TPLANTS.2012.02.003.

18. Gotoh, E., et al. (2018) ‘Palisade cell shape affects the light-induced chloroplast movements and leaf photosynthesis’, Scientific Reports 2018 8:1, 8(1), pp. 1–9. Available at: 10.1038/s41598-018-19896-9.

19. Grieneisen, V.A., et al. (2007) ‘Auxin transport is sufficient to generate a maximum and gradient guiding root growth’, Nature 2007 449:7165, 449(7165), pp. 1008–1013. Available at: 10.1038/nature06215.

20. Hamant, O. and Traas, J. (2010) ‘The mechanics behind plant development’, New Phytologist, 185(2), pp. 369–385. Available at: 10.1111/J.1469-8137.2009.03100.X.

21. Hernández-Verdeja, T., et al. (2022) ‘GENOMES UNCOUPLED1 plays a key role during the de-etiolation process in Arabidopsis’, New Phytologist, 235(1), pp. 188–203. Available at: 10.1111/NPH.18115.

22. Higaki, T., et al. (2016) ‘A Theoretical Model of Jigsaw-Puzzle Pattern Formation by Plant Leaf Epidermal Cells’. Available at: 10.1371/journal.pcbi.1004833.

23. Houbaert, A., et al. (2018) ‘POLAR-guided signalling complex assembly and localization drive asymmetric cell division’, Nature 2018 563:7732, 563(7732), pp. 574–578. Available at: 10.1038/s41586-018-0714-x.

24. Hu, W., et al. (2013) ‘Unanticipated regulatory roles for Arabidopsis phytochromes revealed by null mutant analysis’, Proceedings of the National Academy of Sciences of the United States of America, 110(4), pp. 1542–1547. Available at: 10.1073/PNAS.1221738110.

25. Huang, D.W., et al. (2007) ‘The DAVID Gene Functional Classification Tool: A novel biological module-centric algorithm to functionally analyze large gene lists’, Genome Biology, 8(9), pp. 1–16. Available at: 10.1186/GB-2007-8-9-R183.

26. Hwang, G., et al. (2017) ‘PIF4 promotes expression of LNG1 and LNG2 to induce thermomorphogenic growth in arabidopsis’, Frontiers in Plant Science, 8, p. 282656. Available at: 10.3389/FPLS.2017.01320.

27. Jarvis, P. and López-Juez, E. (2013) ‘Biogenesis and homeostasis of chloroplasts and other plastids’, Nature Reviews Molecular Cell Biology 2013 14:12, 14(12), pp. 787–802. Available at: 10.1038/nrm3702.

28. Jedynak, P., et al. (2022) ‘Dynamics of Etiolation Monitored by Seedling Morphology, Carotenoid Composition, Antioxidant Level, and Photoactivity of Protochlorophyllide in Arabidopsis thaliana’, Frontiers in Plant Science, 12, p. 772727. Available at: 10.3389/FPLS.2021.772727

29. Jiao, Y., Lau, O.S. and Deng, X.W. (2007) ‘Light-regulated transcriptional networks in higher plants’, Nature Reviews Genetics 2007 8:3, 8(3), pp. 217–230. Available at: 10.1038/nrg2049.

30. Jonsson, K., Hamant, O. and Bhalerao, R.P. (2022) ‘Plant cell walls as mechanical signaling hubs for morphogenesis’, Current Biology, 32(7), pp. R334–R340. Available at: 10.1016/j.cub.2022.02.036.

31. Kim, J., et al. (2016) ‘Epidermal Phytochrome B Inhibits Hypocotyl Negative Gravitropism Non-Cell-Autonomously’, The Plant Cell, 28(11), p. 2770. Available at: 10.1105/TPC.16.00487.

32. Kim, Sara, et al. (2020) ‘The epidermis coordinates thermoresponsive growth through the phyB-PIF4-auxin pathway’, Nature Communications 2020 11:1, 11(1), pp. 1–13. Available at: 10.1038/s41467-020-14905-w.

33. Kirchhelle, C., et al. (2019) ‘Two mechanisms regulate directional cell growth in Arabidopsis lateral roots’, eLife, 8. Available at: 10.7554/ELIFE.47988.

34. Labbé, R.M., et al. (2012) ‘A Comparative Transcriptomic Analysis Reveals Conserved Features of Stem Cell Pluripotency in Planarians and Mammals’, Stem Cells, 30(8), pp. 1734–1745. Available at: 10.1002/STEM.1144.

35. Leivar, P., et al. (2008) ‘Multiple Phytochrome-Interacting bHLH Transcription Factors Repress Premature Seedling Photomorphogenesis in Darkness’, Current Biology, 18(23), pp. 1815–1823. Available at: 10.1016/j.cub.2008.10.058.

36. Leivar, P., et al. (2009) ‘Definition of early transcriptional circuitry involved in light-induced reversal of PIF-imposed repression of photomorphogenesis in young Arabidopsis seedlings.’, The Plant cell, 21(11), pp. 3535–53. Available at: 10.1105/tpc.109.070672.

37. Leivar, P. and Monte, E. (2014) ‘PIFs: Systems Integrators in Plant Development’, The Plant Cell, 26(1), pp. 56–78. Available at: 10.1105/TPC.113.120857.

38. Lin, D., Ren, H. and Fu, Y. (2015) ‘ROP GTPase-mediated auxin signaling regulates pavement cell interdigitation in Arabidopsis thaliana’, Journal of Integrative Plant Biology, 57(1), pp. 31–39. Available at: 10.1111/JIPB.12281.

39. Martin, G., et al. (2016) ‘Phytochrome and retrograde signalling pathways converge to antagonistically regulate a light-induced transcriptional network’, Nature Communications 2016 7:1, 7(1), pp.1–10. Available at: 10.1038/ncomms11431.

40. Martín, G., et al. (2020) ‘The photoperiodic response of hypocotyl elongation involves regulation of CDF1 and CDF5 activity’, Physiologia Plantarum, 169(3), pp. 480–490. Available at: 10.1111/PPL.13119.

41. Martín, G., et al. (2021) ‘Alternative splicing landscapes in Arabidopsis thaliana across tissues and stress conditions highlight major functional differences with animals’, Genome Biology, 22(1), pp. 1–26. Available at: 10.1186/S13059-020-02258-Y/FIGURES/6.

42. Martín, G. and Duque, P. (2021) ‘Tailoring photomorphogenic markers to organ growth dynamics’, Plant Physiology, 186(1), pp. 239–249. Available at: 10.1093/plphys/kiab083

43. Monte, E., et al. (2004) ‘The phytochrome-interacting transcription factor, PIF3, acts early, selectively, and positively in light-induced chloroplast development’, Proceedings of the National Academy of Sciences of the United States of America, 101(46), pp. 16091–16098. Available at: 10.1073/PNAS.0407107101

44. Neff, M.M. and Van Volkenburgh, E. (1994) ‘Light-Stimulated Cotyledon Expansion in Arabidopsis Seedlings (The Role of Phytochrome B)’, Plant Physiol Available at: 10.1104/pp.104.3.1027.

45. Ni, M., Tepperman, J.M. and Quail, P.H. (1999) ‘Binding of phytochrome B to its nuclear signalling partner PIF3 is reversibly induced by light’, Nature 1999 400:6746, 400(6746), pp. 781–784. Available at: 10.1038/23500.

46. Outlaw, W.H., Schmuck, C.L. and Tolbert, N.E. (1976) ‘Photosynthetic Carbon Metabolism in the Palisade Parenchyma and Spongy Parenchyma of Vicia faba L’, Plant physiology, 58(2), pp. 186–189. Available at: 10.1104/PP.58.2.186.

47. Park, E., et al. (2012) ‘Phytochrome B inhibits binding of phytochrome-interacting factors to their target promoters’, The Plant Journal, 72(4), pp. 537–546. Available at: 10.1111/J.1365-313X.2012.05114.X.

48. Pfeiffer, A., et al. (2014) ‘Combinatorial Complexity in a Transcriptionally Centered Signaling Hub in Arabidopsis’, Molecular Plant, 7(11), pp. 1598–1618. Available at: 10.1093/MP/SSU087.

49. Procko, C., et al. (2022) ‘Leaf cell-specific and single-cell transcriptional profiling reveals a role for the palisade layer in UV light protection’, The Plant Cell, 34(9), pp. 3261–3279. Available at: 10.1093/PLCELL/KOAC167.

50. Reed, J.W., et al. (1993) ‘Mutations in the gene for the red/far-red light receptor phytochrome B alter cell elongation and physiological responses throughout Arabidopsis development.’, The Plant Cell, 5(2), pp. 147–157. Available at: 10.1105/TPC.5.2.147.

51. Reed, J.W., et al. (1994) ‘Phytochrome A and Phytochrome B Have Overlapping but Distinct Functions in Arabidopsis Development’, Plant Physiology, 104(4), pp. 1139–1149. Available at: 10.1104/PP.104.4.1139.

52. Robinson, S. (2021) ‘Mechanobiology of cell division in plant growth’, New Phytologist, 231(2), pp. 559–564. Available at: 10.1111/NPH.17369.

53. Rovira, A., et al. (2024) ‘PIF transcriptional regulators are required for rhythmic stomatal movements’, Nature communications, 15(1), p. 4540. Available at: 10.1038/S41467-024-48669-4.

54. Ruckle, M.E. and Larkin, R.M. (2009) ‘Plastid signals that affect photomorphogenesis in Arabidopsis thaliana are dependent on GENOMES UNCOUPLED 1 and cryptochrome 1’, New Phytologist, 182(2), pp. 367–379. Available at: 10.1111/j.1469-8137.2008.02729.x.

55. Sapala, A., et al. (2018) ‘Why plants make puzzle cells, and how their shape emerges’, eLife, 7. Available at: 10.7554/ELIFE.32794.

56. Schneider, T., Kaul, C.M. and Pressel, K.G. (2019) ‘Possible climate transitions from breakup of stratocumulus decks under greenhouse warming’, Nature Geoscience 2019 12:3, 12(3), pp. 163–167. Available at: 10.1038/s41561-019-0310-1.

57. Sharma, A., et al. (2023) ‘ELONGATED HYPOCOTYL5 (HY5) and HY5 HOMOLOGUE (HYH) maintain shade avoidance suppression in UV-B’, The Plant journal: for cell and molecular biology, 115(5), pp. 1394–1407. Available at: 10.1111/TPJ.16328.

58. Sheerin, D.J., et al. (2015) ‘Light-Activated Phytochrome A and B Interact with Members of the SPA Family to Promote Photomorphogenesis in Arabidopsis by Reorganizing the COP1/SPA Complex’, The Plant Cell, 27(1), pp. 189–201. Available at: 10.1105/TPC.114.134775.

59. Shi, H., et al. (2018) ‘Genome-wide regulation of light-controlled seedling morphogenesis by three families of transcription factors’, Proceedings of the National Academy of Sciences of the United States of America, 115(25), pp. 6482–6487. Available at: 10.1073/PNAS.1803861115.

60. Shin, J., et al. (2009) ‘Phytochromes promote seedling light responses by inhibiting four negatively-acting phytochrome-interacting factors’, Proceedings of the National Academy of Sciences of the United States of America, 106(18), pp. 7660–7665. Available at: 10.1073/PNAS.0812219106.

61. Shin, J., Park, E. and Choi, G. (2007) ‘PIF3 regulates anthocyanin biosynthesis in an HY5-dependent manner with both factors directly binding anthocyanin biosynthetic gene promoters in Arabidopsis’, The Plant Journal, 49(6), pp. 981–994. Available at: 10.1111/j.1365-313X.2006.03021.x.

62. Smit, M.E., et al. (2023) ‘Extensive embryonic patterning without cellular differentiation primes the plant epidermis for efficient post-embryonic stomatal activities’, Developmental Cell, 58(6), pp. 506–521.e5. Available at: 10.1016/j.devcel.2023.02.014.

63. Stoynova-Bakalova, E., et al. (2004) ‘Cell division and cell expansion in cotyledons of Arabidopsis seedlings’, New Phytologist, 162(2), pp. 471–479. Available at: 10.1111/J.1469-8137.2004.01031.X.

64. Sullivan, J.A. and Gray, J.C. (1999) ‘Plastid translation is required for the expression of nuclear photosynthesis genes in the dark and in roots of the pea lip1 mutant’, The Plant cell, 11(5), pp. 901–910. Available at: 10.1105/TPC.11.5.901.

65. Sun, N., et al. (2016) ‘Arabidopsis saurs are critical for differential light regulation of the development of various organs’, Proceedings of the National Academy of Sciences of the United States of America, 113(21), pp. 6071–6076. Available at: 10.1073/PNAS.1604782113.

66. Tepperman, J.M., Hwang, Y.S. and Quail, P.H. (2006) ‘phyA dominates in transduction of red-light signals to rapidly responding genes at the initiation of Arabidopsis seedling de-etiolation’, The Plant Journal, 48(5), pp. 728–742. Available at: 10.1111/J.1365-313X.2006.02914.X.

67. Ulijasz, A.T., et al. (2010) ‘Structural basis for the photoconversion of a phytochrome to the activated Pfr form’, Nature, 463(7278), pp. 250–254. Available at: 10.1038/NATURE08671.

68. Veciana, N., et al. (2022) ‘BBX16 mediates the repression of seedling photomorphogenesis downstream of the GUN1/GLK1 module during retrograde signalling’, New Phytologist, 234(1), pp. 93–106. Available at: 10.1111/NPH.17975.

69. Verger, S., et al. (2018) ‘A tension-adhesion feedback loop in plant epidermis’, eLife, 7. Available at: 10.7554/ELIFE.34460.

70. Wang, L., et al. (2019) ‘Cotyledons contribute to plant growth and hybrid vigor in Arabidopsis’, Planta, 249(4), pp. 1107–1118. Available at: 10.1007/S00425-018-3068-6.

71. Wang, W., et al. (2021) ‘Direct phosphorylation of HY5 by SPA kinases to regulate photomorphogenesis in Arabidopsis’, New Phytologist, 230(6), pp. 2311–2326. Available at: 10.1111/NPH.17332.

72. Wu, G.-Z., et al. (2018) ‘Control of Retrograde Signaling by Rapid Turnover of GENOMES UNCOUPLED1’, Plant Physiology, 176(3), pp. 2472–2495. Available at: 10.1104/PP.18.00009.

73. Žádníkova, P., et al. (2010) ‘Role of PIN-mediated auxin efflux in apical hook development of Arabidopsis thaliana’, Development, 137(4), pp. 607–617. Available at: 10.1242/DEV.041277.

74. Zhang, X., et al. (2020) ‘AtOFPs regulate cell elongation by modulating microtubule orientation via direct interaction with TONNEAU2’, Plant Science, 292, p. 110405. Available at: 10.1016/J.PLANTSCI.2020.110405.

75. Zhang, Y., et al. (2015) ‘Brassinosteroid is required for sugar promotion of hypocotyl elongation in Arabidopsis in darkness’, Planta, 242(4), pp. 881–893. Available at: 10.1007/S00425-015-2328-Y.

76. Zhang, Z., et al. (2020) ‘A WOX/Auxin Biosynthesis Module Controls Growth to Shape Leaf Form’, Current Biology, 30(24), pp. 4857–4868.e6. Available at: 10.1016/J.CUB.2020.09.037.

77. Zhao, Q.P., et al. (2020) ‘Cryptochrome-mediated hypocotyl phototropism was regulated antagonistically by gibberellic acid and sucrose in Arabidopsis’, Journal of Integrative Plant Biology, 62(5), pp. 614–630. Available at: 10.1111/JIPB.12813.

78. Zhao, X., Huang, J. and Chory, J. (2019) ‘GUN1 interacts with MORF2 to regulate plastid RNA editing during retrograde signaling’, Proceedings of the National Academy of Sciences, 116(20), pp. 10162–10167. Available at: 10.1073/PNAS.1820426116.

