## Supplementary figures and images for "A PIF- and GUN1-regulated switch in cell axis growth drives cotyledon expansion through tissue-specific cell expansion and division"

### Supplemental Figures

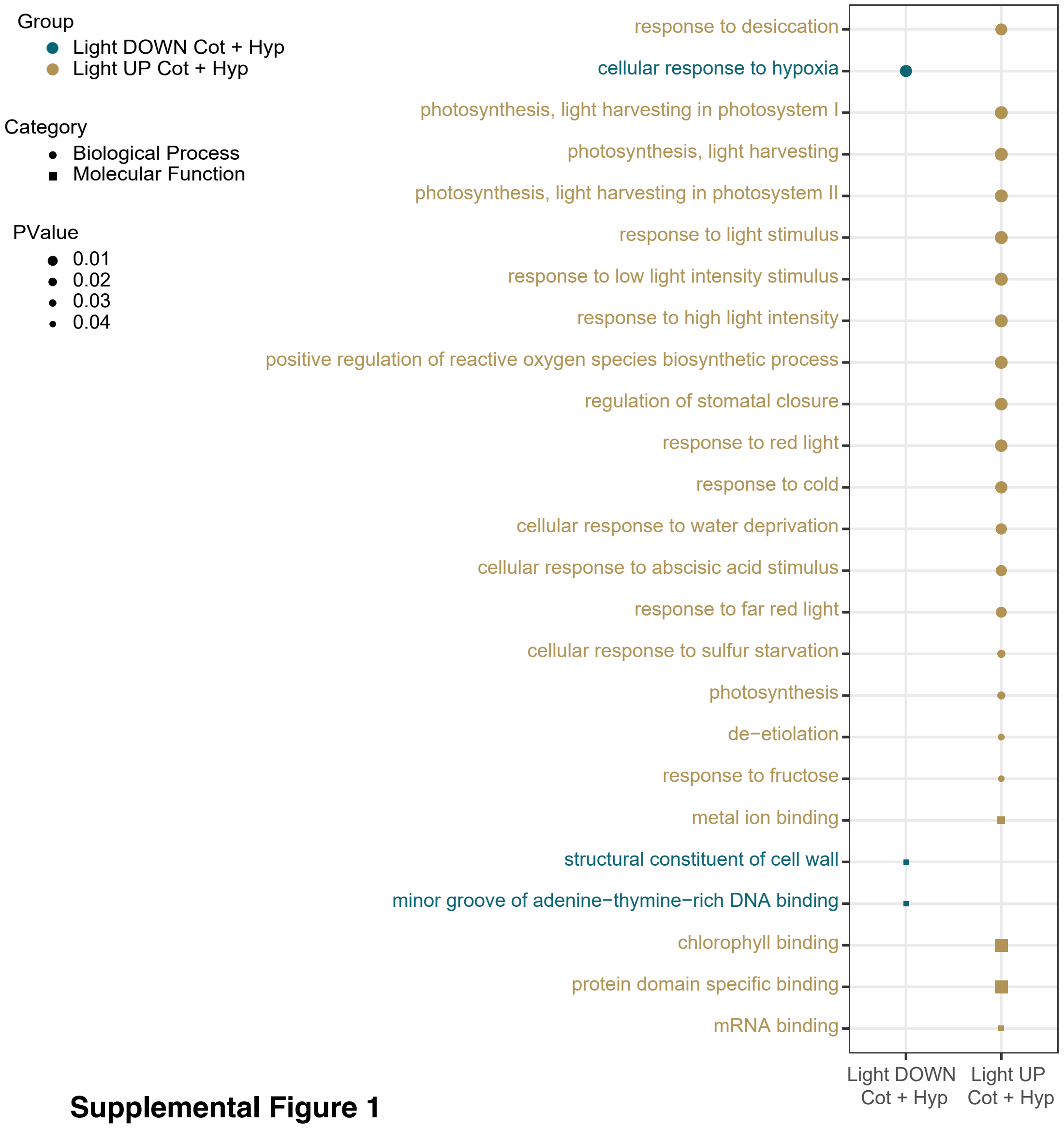


**Supplemental Figure 1**

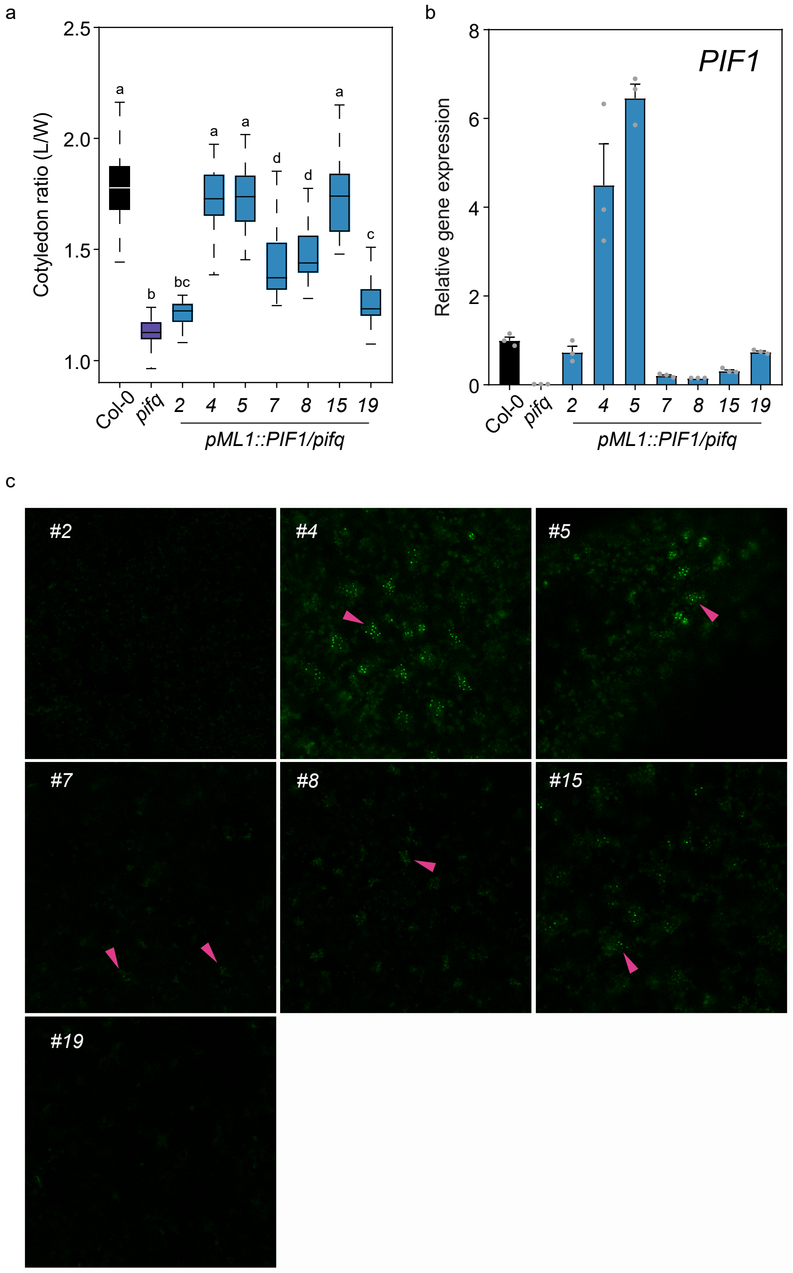


**Supplemental Figure 2**

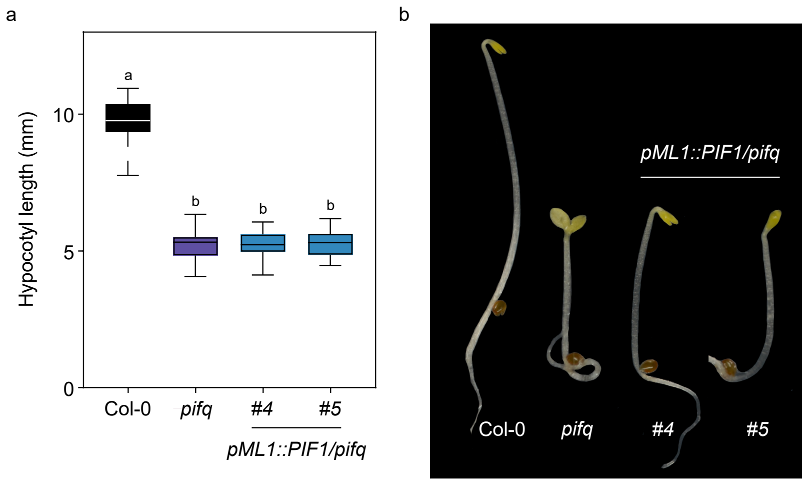


**Supplemental Figure 3**
